# Maize *Brittle Stalk2-Like3,* encoding a COBRA protein, functions in cell wall formation and carbohydrate partitioning

**DOI:** 10.1101/2021.06.04.447139

**Authors:** Benjamin T. Julius, Tyler J. McCubbin, Rachel A. Mertz, Nick Baert, Jan Knoblauch, DeAna G. Grant, Kyle Conner, Saadia Bihmidine, Paul Chomet, Ruth Wagner, Jeff Woessner, Karen Grote, Jeanette Peevers, Thomas L. Slewinski, Maureen C. McCann, Nicholas C. Carpita, Michael Knoblauch, David M. Braun

**Affiliations:** Divisions of Plant and Biological Sciences, Interdisciplinary Plant Group, and the Missouri Maize Center, University of Missouri, Columbia, MO 65211 USA; School of Biological Sciences, Washington State University, Pullman, Washington 99164, USA; NRGene Inc., 8910 University Center Lane, Suite 400, San Diego, CA 92122 USA; Bayer Crop Science, Chesterfield, MO 63017 USA; Department of Biological Sciences, Purdue University, West Lafayette, IN 47907 USA; Department of Botany and Plant Pathology, Purdue University, West Lafayette, IN 47907 USA; Purdue Center for Plant Biology, Purdue University, West Lafayette, IN 47907 USA; Electron Microscopy Core Facility, University of Missouri, W136 Veterinary Medicine Building 1600 East Rollins Street, Columbia, MO 65211, USA

**Keywords:** Carbohydrate partitioning, cell wall, cellulose, *cobra*, *cpd28*, phloem, sucrose

## Abstract

Carbohydrate partitioning from leaves to sink tissues is essential for plant growth and development. The maize (*Zea mays*) recessive *carbohydrate partitioning defective28* (*cpd28*) and *cpd47* mutants exhibit leaf chlorosis and accumulation of starch and soluble sugars. Transport studies with ^14^C-sucrose (Suc) found drastically decreased export from mature leaves in *cpd28* and *cpd47* mutants relative to wild-type siblings. Consistent with decreased Suc export, *cpd28* mutants exhibited decreased phloem pressure in mature leaves, and altered phloem cell wall ultrastructure in immature and mature leaves. We identified the causative mutations in the *Brittle Stalk2-Like3* (*BK2L3*) gene, a member of the COBRA family, which is involved in cell wall development across angiosperms. None of the previously characterized *COBRA* genes are reported to affect carbohydrate export. Consistent with other characterized COBRA members, the BK2L3 protein localized to the plasma membrane, and the mutants condition a dwarf phenotype in dark-grown shoots and primary roots, as well as the loss of anisotropic cell elongation in the root elongation zone. Likewise, both mutants exhibit a significant cellulose deficiency in mature leaves. Therefore, *BK2L3* functions in tissue growth and cell wall development, and this work elucidates a unique connection between cellulose deposition in the phloem and whole-plant carbohydrate partitioning.

**Funding Information:** The research was supported by US National Science Foundation Plant Genome Research Program grants (IOS-1025976 and IOS-1444448) to DMB.

## INTRODUCTION

Many products used by society, from clothing to food to alternative energy sources, are harvested from various crops and vegetation. These resources are the result of carbohydrate partitioning, the transport of carbohydrates synthesized in photosynthetic source tissues (i.e., leaves) to various non-photosynthetic sink tissues (e.g., roots, reproductive organs, stem) for proper growth, development, and yield (Lalonde et al., 2003; Braun et al., 2014; Julius et al., 2017). In maize (*Zea mays*) and other crops, sucrose (Suc) is synthesized in the mesophyll cells of source leaves and transported to the phloem, the vascular tissue responsible for long-distance transport of assimilates (van Bel, 2003; Braun and Slewinski, 2009; Ayre, 2011). The phloem is primarily composed of three cell types: phloem parenchyma cells, companion cells and sieve elements. The latter two cell types are found in complexes as the sieve elements are enucleate and rely on associated companion cells for metabolic support (Esau, 1977; van Bel and Knoblauch, 2000). Suc is transported from the mesophyll to the bundle sheath cells and on into the phloem parenchyma cells via cell wall-spanning cytoplasmic connections called plasmodesmata (Evert, 1982). In maize, plasmodesmata are rare at the interface of the bundle sheath or phloem parenchyma cells and companion cell-sieve element complex (Evert et al., 1978). Therefore, Suc is effluxed into the apoplastic cell wall space from these cell types by SWEET proteins and imported into the companion cell by the SUCROSE TRANSPORTER1 protein (Slewinski et al., 2009; Baker et al., 2016; Bezrutczyk et al., 2018; Bezrutczyk et al., 2021). Suc undergoes symplastic transport via plasmodesmata from the companion cell into the sieve element (Evert, 1982).

In grass leaves, there are two types of sieve elements, thin-walled and thick-walled, that have distinct functions in long-distance assimilate transport and can be distinguished based on their position within a vein and the thickness of their cell walls (Botha, 2005). In plants, sieve elements are connected end-to-end by porous sieve plates forming a sieve tube, a transport path throughout the plant allowing for assimilate export from source leaf tissue to various sink organs (Esau, 1977; Evert, 1982). Suc, potassium, chloride, amino acids, and other compounds are translocated from source to sink tissues in the phloem sap and generate a very high osmolarity (>1.6-2.3 M) (Ohshima et al., 1990; Weiner et al., 1991). The high Suc and other osmolyte concentrations in the source leaf phloem attract water from the nearby xylem to create a high hydrostatic pressure (Münch, 1930). Conversely, Suc unloading from the phloem in sink tissues decreases the osmolyte concentration and results in lower hydrostatic pressure. The difference in pressure between source and sink tissues drives bulk flow through the phloem (van Bel, 2003).

Several genes in maize have been characterized to influence carbohydrate partitioning, and when mutated exhibit a suite of phenotypes such as decreased plant growth, chlorotic leaves, and reduced fertility, all associated with hyperaccumulation of carbohydrates in leaves. The mutants can be classified into three groups: 1) *psychedelic* (*psc*), *tie-dyed1* (*tdy1*), *tdy2,* and *carbohydrate partitioning defective33* (*cpd33*), in which the causes of inhibited carbohydrate transport are unknown, although the *Tdy* and *Cpd33* loci are hypothesized to function in companion cell-sieve element plasmodesmata transport (Baker and Braun, 2008; Ma et al., 2008; Ma et al., 2009; Slewinski and Braun, 2010; Slewinski et al., 2012; Baker et al., 2013; Tran et al., 2019), 2) *sucrose transporter1* (*sut1*), *sut2*, and the triple mutant *sweet13a*, *b*, and *c,* which function as Suc transporters (Carpaneto et al., 2005; Slewinski et al., 2010; Baker et al., 2016; Leach et al., 2017; Bezrutczyk et al., 2018), and 3) *sucrose export defective1* (*sxd1*) and *Carbohydrate partitioning defective1* (*Cpd1*), which result in ectopic callose deposits that block Suc movement at various points along its transport pathway (Russin et al., 1996; Julius et al., 2018). These mutants are all impaired in their ability to export carbohydrates from their source tissue resulting in their respective shared phenotypes, despite different underlying mechanistic defects among the three groups of mutants.

A major part of the Suc imported into growing sink tissues is converted into carbon skeletons that provide the building materials for new cells, including the cell wall, a heterogeneous polysaccharide matrix surrounding the cell (Ruan, 2014; Liepman et al., 2018; Rui and Dinneny, 2020). Additionally, significant differences have been found between the composition of cell walls of eudicots, non-commelinoid monocots, and gymnosperms (Type I) compared with grasses (Type II) (Carpita and Gibeaut, 1993). Both Type I and Type II cell walls contain cellulose; however, in Type I walls, cellulose is found in a network of mostly xyloglucan, pectin and structural proteins, whereas in Type II walls, the cellulose is networked with mostly glucuronoarabinoxylans and low levels of pectin and structural proteins. Cellulose is a major component of both primary cell walls, which control the anisotropic growth of cells, and secondary cell walls, which are deposited upon the completion of cell elongation and are crucial for structural support of the plant tissue (Carpita, 1996). Cellulose is synthesized at the plasma membrane by cellulose synthase ‘rosette’ complexes (Giddings Jr et al., 1980; Mueller and Brown Jr, 1980). These complexes are composed of several distinct cellulose synthase isoforms, sub-functionalized for the deposition of cellulose in primary or secondary cell walls (Taylor et al., 2003; Appenzeller et al., 2004).

Additional proteins are required for proper cellulose synthesis and orientation, such as the COBRA proteins (Schindelman et al., 2001). The COBRA family was first identified in *Arabidopsis thaliana* with the characterization of the *cobra* mutant that exhibited abnormal root growth (Benfey et al., 1993). Several additional gene family members have been characterized in Arabidopsis, maize, rice (*Oryza sativa*), cotton (*Gossypium hirsutum*), and tomato (*Solanum lycopersicum*), affecting plant height, leaf and stem brittleness, and the development of root hairs, seed coat, pollen, and cotton fiber cells (Hochholdinger et al., 2008; Cao et al., 2012; Li et al., 2013; Liu et al., 2013; Ben-Tov et al., 2015; Niu et al., 2015). For example, *Brittle Stalk2 (Bk2)* was the first COBRA family member to be identified in maize (Ching et al., 2006; Sindhu et al., 2007), and was proposed to function in lignin-cellulose patterning required for proper secondary cell wall development and tissue flexibility at maturity (Sindhu et al., 2007). In most cases, COBRA gene family mutants exhibited decreased cellulose content (Roudier et al., 2002; Li et al., 2003; Brown et al., 2005; Sindhu et al., 2007; Dai et al., 2011), while overexpression of COBRA in tomato exhibited increased cellulose levels (Cao et al., 2012). COBRA proteins contain four conserved domains: an N-terminal protein targeting domain, a carbohydrate-binding motif, a conserved CCVS domain of unknown function, and a hydrophobic C-terminal tail domain. Both the N-terminal domain and the hydrophobic C-terminal domain are cleaved as post-translational modifications, and a glycosylphosphatidylinositol (GPI) anchor is attached at the C-terminal end of the protein after cleavage at the ω-residue (Roudier et al., 2002; Brady et al., 2007). The GPI anchor integrates into the plasma membrane with the remaining peptide sequence located in the apoplast (Eisenhaber et al., 2003). The carbohydrate-binding motif binds cellulose, and disruption of this domain inhibits proper function of COBRA family members (Sato et al., 2010). To date, COBRA mutants have no reported carbohydrate export defects.

In this paper we determined that the allelic *carbohydrate partitioning defective28* (*cpd28*) and *cpd47* mutations occur in the maize *Brittle stalk2-like3 (BK2L3)* gene. *BK2L3* is a COBRA gene and functions in cellulose deposition, affecting the ultrastructure of the sieve element cell wall, and carbohydrate partitioning. This research demonstrates the physiological importance of proper phloem cell wall formation on whole-plant carbohydrate partitioning.

## RESULTS

### *cpd28* and *cpd47* mutants are dwarfed and exhibit reduced photosynthesis

The *cpd28* and *cpd47* mutants were identified from an EMS-mutagenized population based on a suite of phenotypes, including reduced plant height, chlorosis, and anthocyanin accumulation in the leaves (Figure 1 and Supp. Figure S1). Initially, as leaves develop and emerge from the whorl they appear healthy and normal, with the chlorosis and anthocyanin accumulation developing in a basipetal pattern with continuing exposure to sunlight (Figure 1B, C and Supp. Figure S1B, C). Based on Mendelian segregation ratios both *cpd28* and *cpd47* were determined to be recessive mutants segregating in a 1:3 ratio for mutant to wild-type plants. Due to the similarity in phenotype between the two mutants, heterozygous plants carrying either allele were crossed together, and the resulting offspring segregated the phenotype in a 1:3 mutant to wild-type ratio, supporting that the two mutations are allelic.

**Figure 1.**
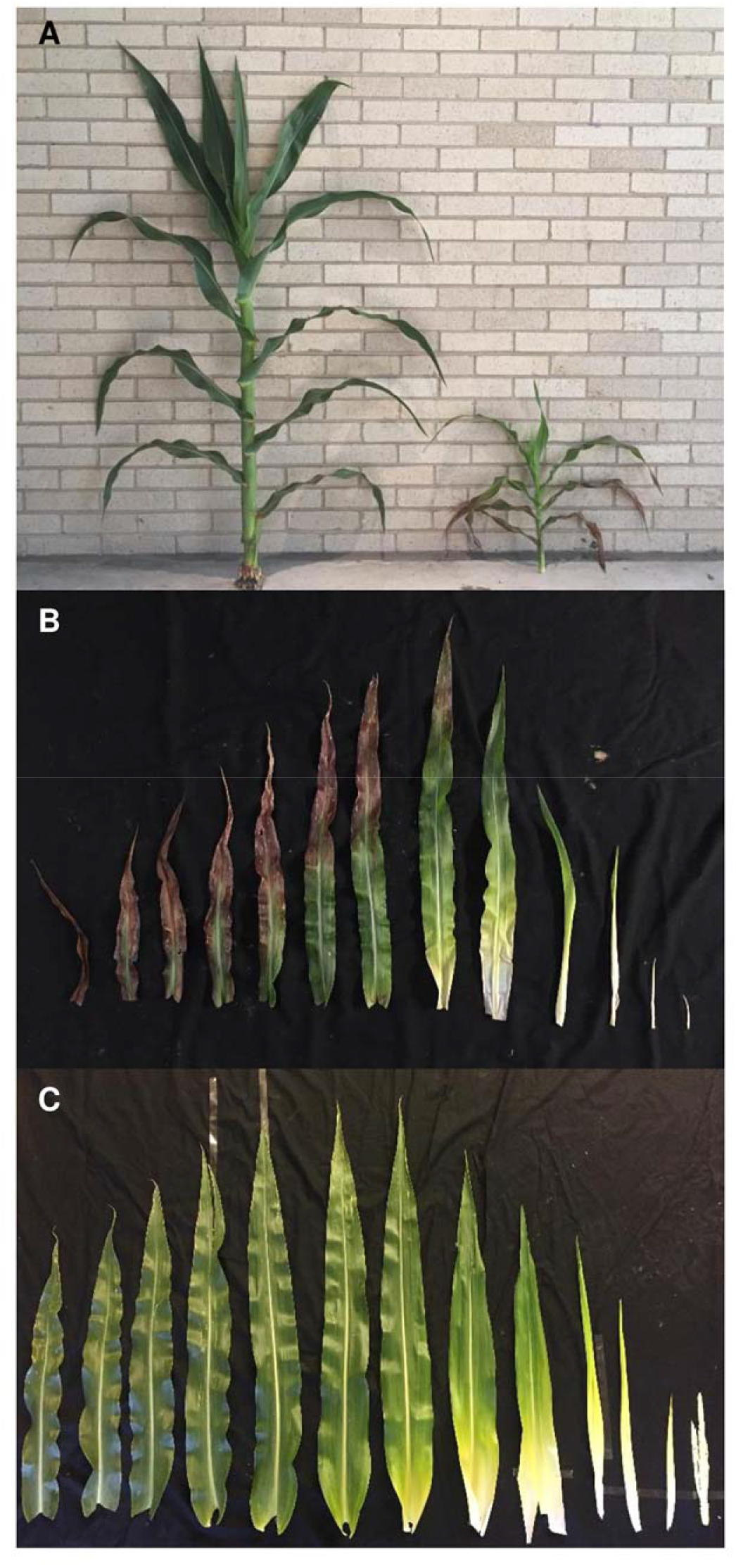
*cpd28* mutant plants are dwarfed relative to wild-type (WT) siblings and exhibit basipetal accumulation of anthocyanin in their leaves. (A) Field grown WT (left) and *cpd28* mutant (right) plants. (B-C) A gradient of *cpd28* (B) and WT (C) leaves from oldest (left) to youngest (right). The mutant leaves accumulate anthocyanins in a basipetal fashion when exposed to sunlight.

Because of the chlorosis in mutant leaves, measurements were taken to determine the differences in photosynthetic parameters between the wild-type and mutant siblings. Both the photosynthetic rate and stomatal conductance were dramatically decreased in the *cpd28* and *cpd47* individuals relative to wild type (Supp. Figure S2).

### *cpd28* and *cpd47* mutants hyperaccumulate carbohydrates in their leaves

The decreased plant height, leaf chlorosis, anthocyanin accumulation, and reduced gas exchange in the *cpd28* and *cpd47* mutants were similar to phenotypes observed in other mutants exhibiting defects in carbohydrate export from their leaves (see Introduction). Therefore, mature *cpd28* and *cpd47* mutants, and their respective wild-type sibling leaves were harvested at end of night, cleared of photosynthetic pigments, and stained with iodine potassium iodide (IKI) for the presence of starch. Both mutants exhibited hyperaccumulation of starch relative to their wild-type siblings (Figure 2A, B and Supp. Figure S3A, B).

**Figure 2.**
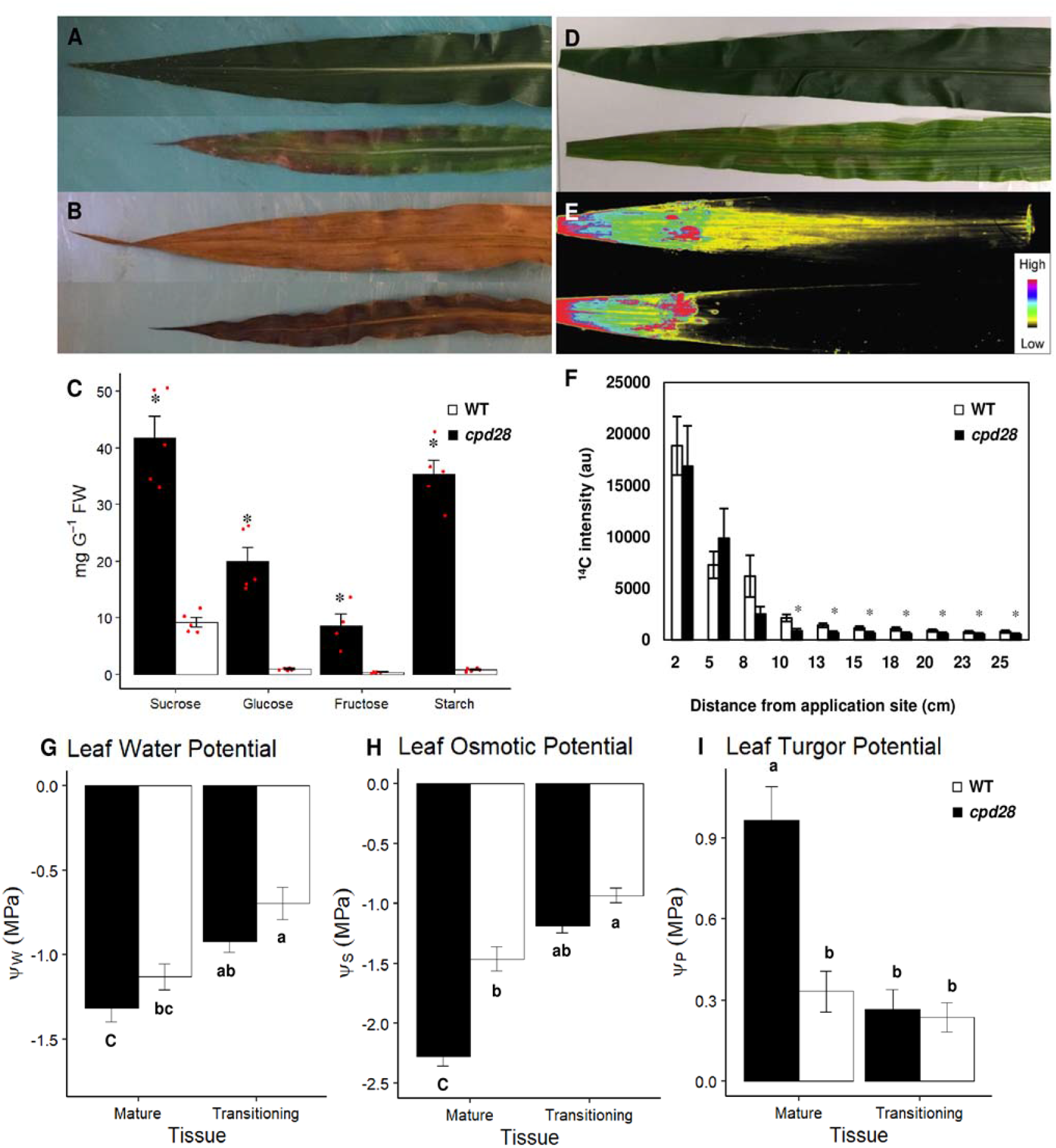
Reduced Suc export leads to hyperaccumulation of starch and soluble sugars in mature *cpd28* mutant leaves. (A-B) Wild-type (WT, top) sibling and *cpd28* mutant (bottom) mature leaves before (A) and after (B) IKI stain. (C) Soluble sugar and starch quantification in WT and *cpd28* mature leaves. (D-F) Export of ^14^C-Suc is reduced in *cpd28* mutant source leaves. (D) Dried WT, top, and *cpd28* mutant leaves, bottom. (E) Autoradiograph showing ^14^C-Suc distribution in WT, top, and *cpd28,* bottom leaves. (F) Quantification of ^14^C-Suc distribution in WT (white bars) and *cpd28* mutant (black bars) leaves. Translocation was measured according to signal intensity at each distance (in cm) from the application site. (G – I) Fully mature mutant leaves, but not mutant leaves still transitioning from sink to source, exhibited significantly increased osmotic potential compared to WT (H), while water potential was not significantly altered (G). Turgor pressure was calculated from these values (I). Error bars are mean ± StdEr for all graphs, red points indicate individual measurements comprising the mean (C). *, P ≤ 0.05 using Student’s *t*-test (C, F); for G – I statistical significance was determined by one-way ANOVA (P≤0.01) with groups (letters) determined by Tukey-Kramer post hoc.

Additionally, quantitative measurements showed that *cpd28* and *cpd47* mature leaves exhibited significantly increased levels of starch, Suc, glucose (Glc), and fructose (Fru) relative to wild-type samples (Figure 2C and Supp. Figure S3C). To understand if the increased soluble sugars and starch in *cpd28* mutant leaves impacted water relations of the plant, we measured the water potential (Ψ_W_) and the osmotic potential (Ψ_S_) of mature leaves, and in leaves transitioning from sink to source tissue of both *cpd28* mutants and wild-type siblings. While neither mature nor transitioning leaves of *cpd28* mutants exhibited a statistically different Ψ_W_ from their respective wild-type sibling leaves (Figure 2G), Ψ_S_ of *cpd28* mature leaves was enhanced by 55% (Figure 2H, -2.28 MPa in *cpd28* mutants compared to -1.46 MPa in wild type). This change in Ψ_S_, coupled with the lack of significant alteration of Ψ_W_, resulted in an over two-fold increase of turgor pressure (Ψ_P_) in *cpd28* mature leaves (Figure 2I). Taken together, these results demonstrated that *cpd28* and *cpd47* mutants have increased carbohydrate accumulation in their leaves, which we hypothesized was due to decreased Suc export from source leaves.

### Mutant leaves exhibit decreased Suc export

To assess long-distance Suc export from leaves, ^14^C-labeled Suc was applied to recently matured source leaves on 5-week-old plants (*cpd28* and *cpd47* mutants and their wild-type respective siblings) and allowed to translocate for 1 hr. We performed the experiment using the most recently matured, fully expanded source leaf of each plant, which was prior to the appearance of chlorosis and anthocyanin accumulation. The radiolabel was transported to the base of the mature wild-type leaf (Figure 2D, E and Supp. Figure S3D, E). However, the transport of radiolabel was severely limited in *cpd28* and *cpd47* plants, and no signal was seen in the basal half of the leaves. When the radioactivity in the images was quantified, a significant reduction in the amount of radiolabel was detected basipetally beginning 10-12 cm from the application site in each mutant compared to wild-type siblings (Figure 2F and Supp. Figure S3F). These data indicate that the hyperaccumulation of carbohydrates in the *cpd28* and *cpd47* mature leaves is a result of decreased Suc export.

### Carbohydrate hyperaccumulation is associated with ectopic callose deposits in phloem of strongly expressing mutant leaves, but not etiolated or transitioning leaves

Veins of *cpd28* and *cpd47* mutants and wild-type sibling leaves were analyzed to determine if there were any abnormalities in regards to anatomy or ectopic deposition of callose potentially causing the inhibited transport of Suc as seen in the *Cpd1* mutant (Julius et al., 2018). Cross sections from the tip of mature leaves displaying anthocyanins (strong and pronounced phenotypic expression) of *cpd28* and *cpd47* mutants were stained with aniline blue and showed callose deposition in the phloem of both mutants but not in wild-type siblings (Supp. Figure S4A-F).

To address whether these deposits might be the cause of the decreased Suc export in the mature leaves, *cpd28, cpd47*, and their wild-type siblings were grown for two weeks under etiolated conditions, which precludes photosynthesis, carbon assimilation, and hence carbohydrate hyperaccumulation in the leaves. Cross-sections stained with aniline blue exhibited no ectopic callose deposits in the phloem of etiolated tissue, or that of light-grown non-photosynthetic immature leaves (Supp. Figure S4G-L). Therefore, we suggest that the ectopic callose deposition observed in the mature phloem of the two mutants grown in field conditions are likely downstream effects of the hyperaccumulation of carbohydrates, and not the cause of the increased carbohydrate levels.

### *cpd28* and *cpd47* exhibit allelic differences in regards to growth

The *cpd28* etiolated seedlings exhibited a severe reduction in growth compared to *cpd47* and their wild-type siblings (Figure 3A, B). Shoots and roots of both mutants were significantly shorter than wild type (Figure 3C, D). The reductions in root growth of the etiolated *cpd28* plants were particularly striking compared to those of wild type or *cpd47* (Figure 3A, B). Consistent with the etiolated shoot data, *cpd28* mutant plants had an average root length ∼6 cm shorter than their wild-type siblings, and *cpd47* plants had an average root length ∼2.5 cm shorter than their wild-type siblings (Figure 3D). Therefore, both *cpd28* and *cpd47* mutants have growth defects in addition to their carbohydrate partitioning phenotype, with *cpd28* being the more severe allele. To determine if the reduced growth was due to decreased carbon mobilization from kernel carbohydrate reserves, we grew seedlings on media supplemented with Suc. The addition of 2% Suc to half-strength Murashige and Skoog (MS) media did not increase root growth of *cpd28* mutants, indicating that carbohydrate availability is not limiting growth in the mutants (Supp. Figure S5).

**Figure 3.**
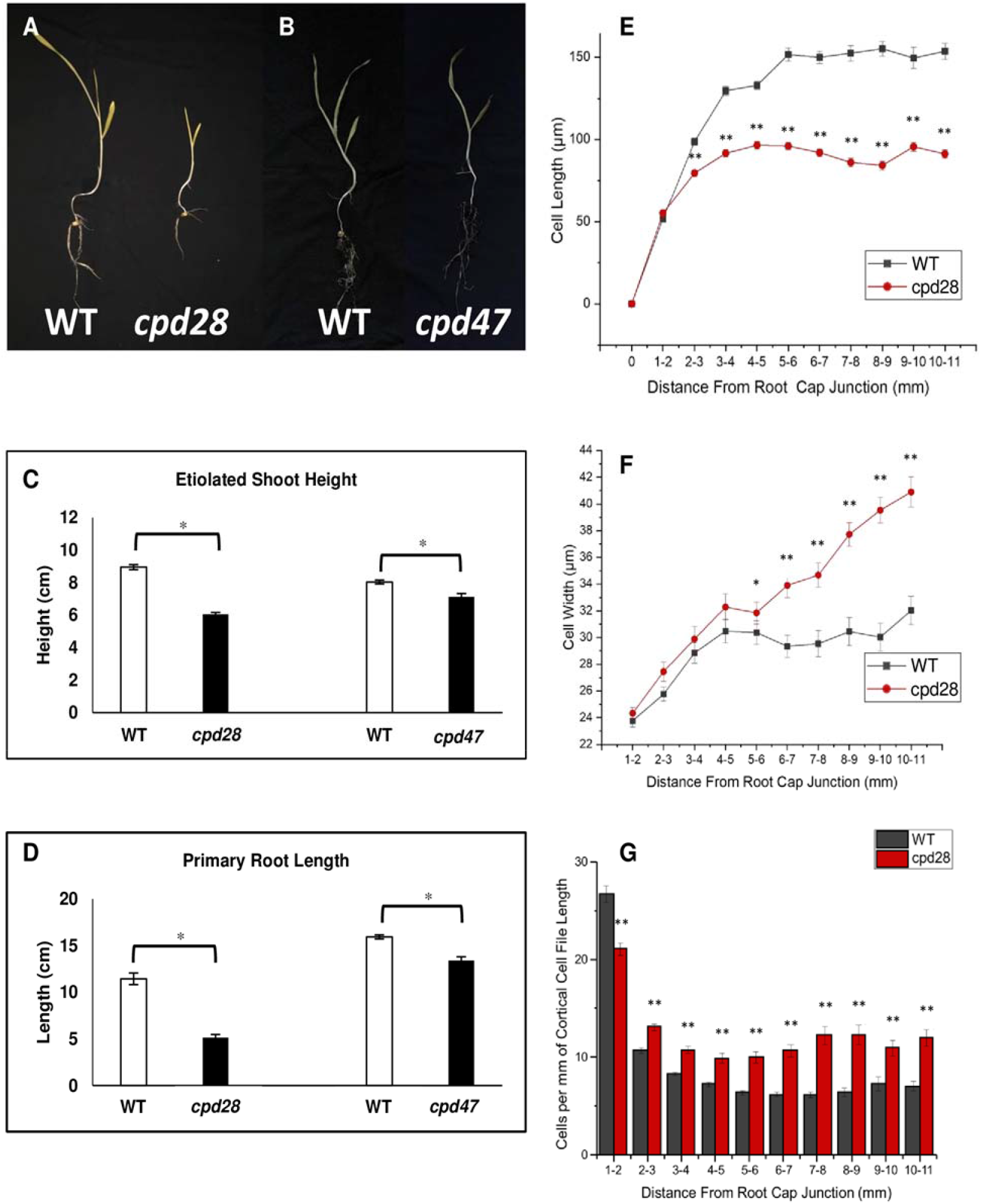
*cpd28* and *cpd47* mutants exhibit dwarf phenotypes and a reduction in cell elongation under etiolated conditions compared to their respective wild-type (WT) siblings. (A-B) Two-week-old etiolated WT sibling (A, left) and *cpd28* mutant (A, right), or WT sibling (B, left) and *cpd47* mutant (B, right) seedlings. (C) Height measurements for *cpd28* and *cpd47* mutants and their respective WT sibling shoots. (D) Length measurements for *cpd28* and *cpd47* mutants and respective WT sibling primary roots. (E) Reduced cell length and increased width (F) in the primary root growth zone. (G) Number of cortical cells in the primary root growth zone. Error bars are mean ± StdEr. * (P ≤ 0.05), ** (P ≤ 0.001) using Student’s *t*-test.

To determine the cause of the decreased root length relative to wild-type siblings, cortical cell length and width measurements were performed in 1 mm intervals of primary root tip median longitudinal sections from *cpd28* mutants and wild-type siblings. There was no difference in cell length and width profiles within the first 2 mm, consisting of the meristematic zone of the primary root (Sharp et al., 1988). However, *cpd28* cells were shorter and wider than the cells observed in wild-type siblings at every site distal to >2 mm from the root cap in the primary root (Figure 3E, F). The inhibition in anisotropic cell elongation begins where cells transition from the meristematic zone to the elongation zone in the root tissue (Sharp et al., 1988). In addition to decreased cell length, there was a significant decrease in cell number in the first 1 mm of *cpd28* mutant root apices compared to wild type (Figure 3G), which also contributes to the shorter roots in the mutant.

To investigate if a similar growth retardation might underlie the reduced shoot growth phenotype, we measured mesophyll cell size in immature, transitioning, and fully mature leaves. No difference in mesophyll cell size was observed between wild type and *cpd28* mutants in immature leaves, but *cpd28* mesophyll cells were 30-40% smaller than in the wild type in the transitioning sink and mature leaves (Supp. Figure S6).

### cpd28 and cpd47 mutations occur in Brittle Stalk2-Like3

To understand the molecular mechanism for the inhibited carbohydrate transport and dwarf phenotypes, we employed a Bulked Segregant Analysis (BSA) to map the mutations to the tip of the long arm of chromosome one. The region was further narrowed with polymorphic PCR markers to a ∼750 kb region, containing 17 predicted protein-encoding genes (Supp. Table S1). To identify the causative mutations for each allele, a whole genome sequencing approach was utilized with DNA isolated from independent pools of >40 mutant plants for each allele. DNA sequences of *cpd28* and *cpd47* were aligned with the B73 reference genome sequence in the region of interest to identify any mutations in the 17 genes. A comparison of polymorphisms between the *cpd28* and the *cpd47* sequences resulted in the identification of a single gene, *Brittle Stalk2-like3* (*BK2L3*), with unique mutations in both.

*BK2L3* is one of nine members of the maize COBRA gene family (Brady et al., 2007). The *cpd28* mutation is a single nucleotide polymorphism resulting in a non-synonymous (missense) amino acid substitution in the first amino acid, changing it from Met to Ile, and therefore resulting in the loss of the start codon, with the next start codon located 30 amino acids downstream and out of frame, resulting in a non-functional protein, if translated (Figure 4A). The *cpd47* mutation is a base pair change, resulting in a missense mutation in amino acid 103, changing a conserved Asp to Asn creating a novel, predicted *N*-glycosylation site at the end of the carbohydrate binding motif (Figure 4B, Supp. Figure S7). Therefore, we conclude that the *cpd28* allele is likely a null allele, and *cpd47* is a hypomorphic allele.

**Figure 4.**
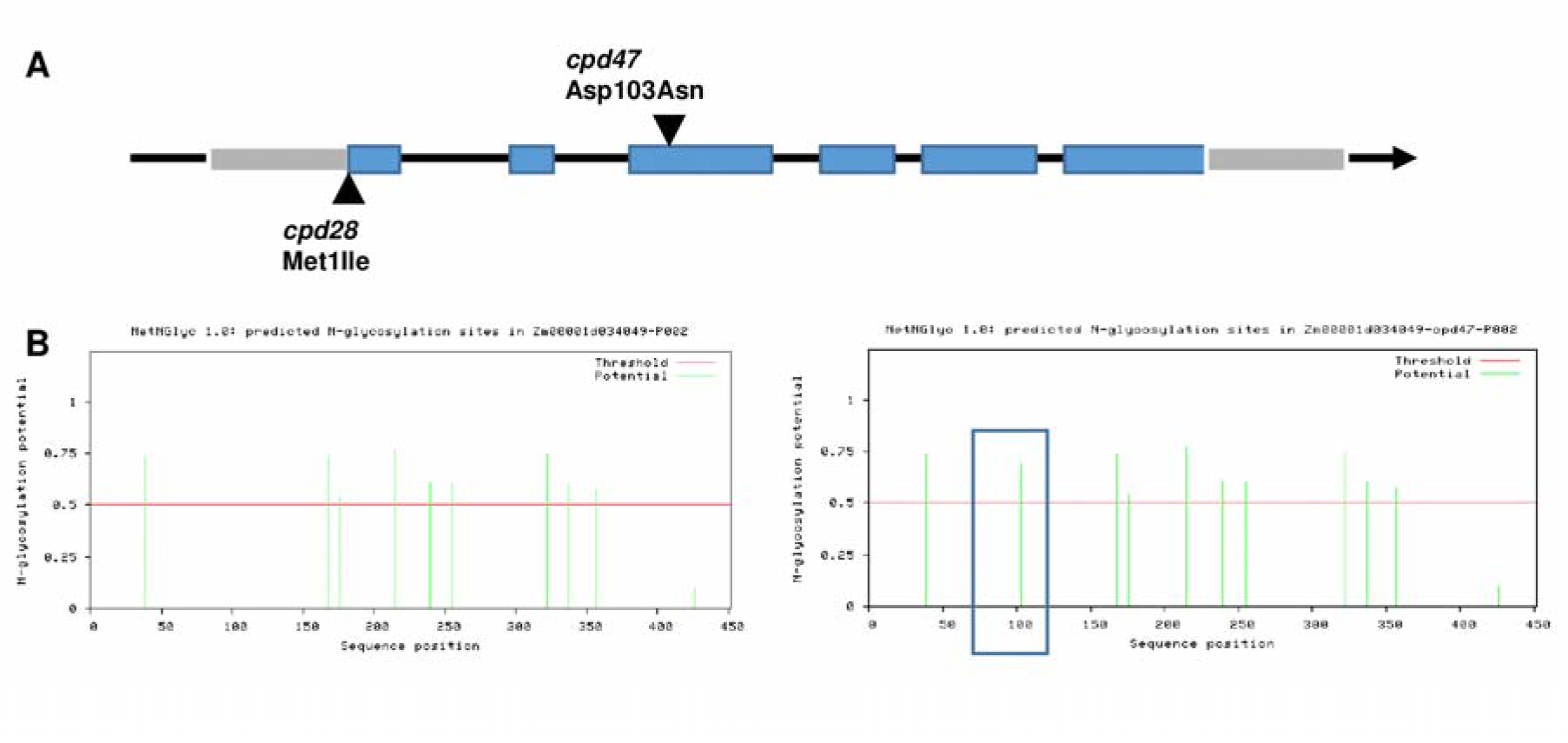
The causative mutations for *cpd28* and *cpd47* were identified in the *Brittle Stalk 2-like3* gene. (A) The *cpd28* mutation results in loss of the start codon. The *cpd47* mutation results in a missense mutation changing an Aspartic Acid (Asp) to Asparagine (Asn). (B) The NetNGlyc 1.0 program was used to predict N-glycosylation sites in the wild-type (left) and *cpd47* mutant (right) BK2L3 protein sequences. The *cpd47* mutation results in an additional predicted N-glycosylation site (blue box).

### BK2L3 localizes to the plasma membrane

Previous studies localized COBRA proteins to the plasma membrane, anchored there via a GPI attachment (Roudier et al., 2002; Dai et al., 2011). To determine whether BK2L3 also localized to the plasma membrane, the yellow fluorescent protein Citrine was translationally inserted immediately after the predicted N-terminal secretory peptide in the BK2L3 protein, upstream of, and in-frame with, the remaining peptide sequence. *Nicotiana benthamiana* leaves were co-infiltrated with 35Spro:BK2L3-Citrine (Figure 5A) and 35Spro:PIP2a-CFP, a tagged aquaporin protein that localizes to the plasma membrane (Figure 5B) (Nelson et al., 2007). The fluorescent signals of the BK2L3-Citrine and PIP2a-CFP proteins overlapped, supporting that BK2L3 localizes to the plasma membrane (Figure 5D).

**Figure 5.**
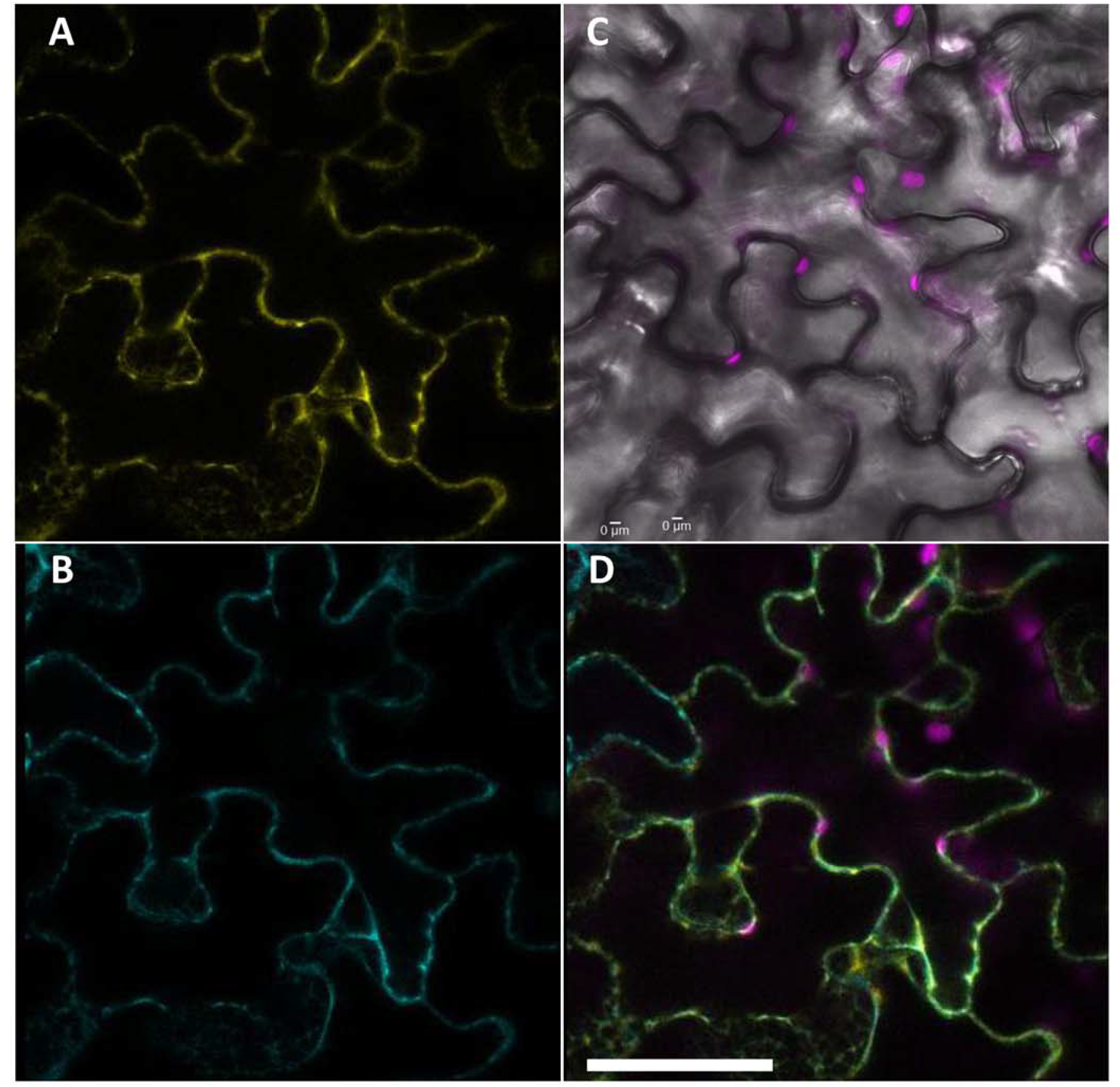
BK2L3 localizes to the plasma membrane. (A-B) Localization of 35Spro:ZmBK2L3-Citrine (A) and the plasma membrane-localized 35Spro:AtPIP2a-CFP (B) in *Nicotiana benthamiana* leaf epidermal cells. (C) Bright-field image of *N. benthamiana* epidermal cells with chloroplasts autofluoresence collected (magenta). (D) An overlay of 35Spro:ZmBK2L3-Citrine, 35Spro:AtPIP2a-CFP, and chloroplast channels. Scale bar = 50 µm.

### *BK2L3* is closely related to *AtCOBRA* and *OsBC1L4*

Several members of the COBRA family have been characterized in Arabidopsis, cotton, tomato, rice, and maize. Phylogenetic analysis separates the protein family into five clades based upon the length of peptide sequences and function. COBRA proteins in clades I, II, and III typically consist of ∼400-450 amino acids, while members in clades IV and V consist of ∼675 amino acids (Figure 6). Clade I includes rice BRITTLE CULM1-LIKE4 (OsBC1L4) and AtCOBRA, which function in the development of primary cell walls, and each exhibits a dwarf phenotype when mutated (Schindelman et al., 2001; Roudier et al., 2002; Zhao et al., 2009; Dai et al., 2011). Similarly, ZmBK2L3 is located in clade I, and etiolated *cpd28* and *cpd47* mutant seedlings also exhibit diminished growth phenotypes (Figure 3). Interestingly, *AtCOBRA*, *OsBC1L4*, and *ZmBK2L3* are co-expressed with primary cell wall cellulose synthases (Dhugga, 2001; Appenzeller et al., 2004; Brady et al., 2007). Clade II contains the COBRA proteins OsBC1, ZmBK2, and AtCOBL4, which have all been characterized as playing important roles in secondary cell wall development (Li et al., 2003; Brown et al., 2005; Sindhu et al., 2007; Liu et al., 2013). Non-structural carbohydrate levels were quantified in the maize *bk2* mutant leaves, but no differences were found in soluble sugars relative to wild type, although starch levels were decreased (Supp. Figure S8). *BK2L3* shows peak relative RNA expression early in organ development, particularly in elongating tissues compared with *Bk2*, which is expressed later in tissue development during secondary cell wall formation (Supp. Figure S9) (Brady et al., 2007; Walley et al., 2016). *BK2L3* is broadly expressed in numerous tissues throughout the plant, including primary roots, lateral roots, lateral root tip meristems, nodal root meristems, young leaves, elongating internodes, pulvinus, developing tassels, anthers, immature ears, expanding silks, developing embryos and endosperms, and maternal pericarp tissues, and it is often expressed at the highest level among maize COBRA-family members (Brady et al., 2007).This broad pattern of gene expression is consistent with a role for BK2L3 in primary cell wall biosynthesis in multiple tissues (Li et al., 2010; Wang et al., 2014).

**Figure 6.**
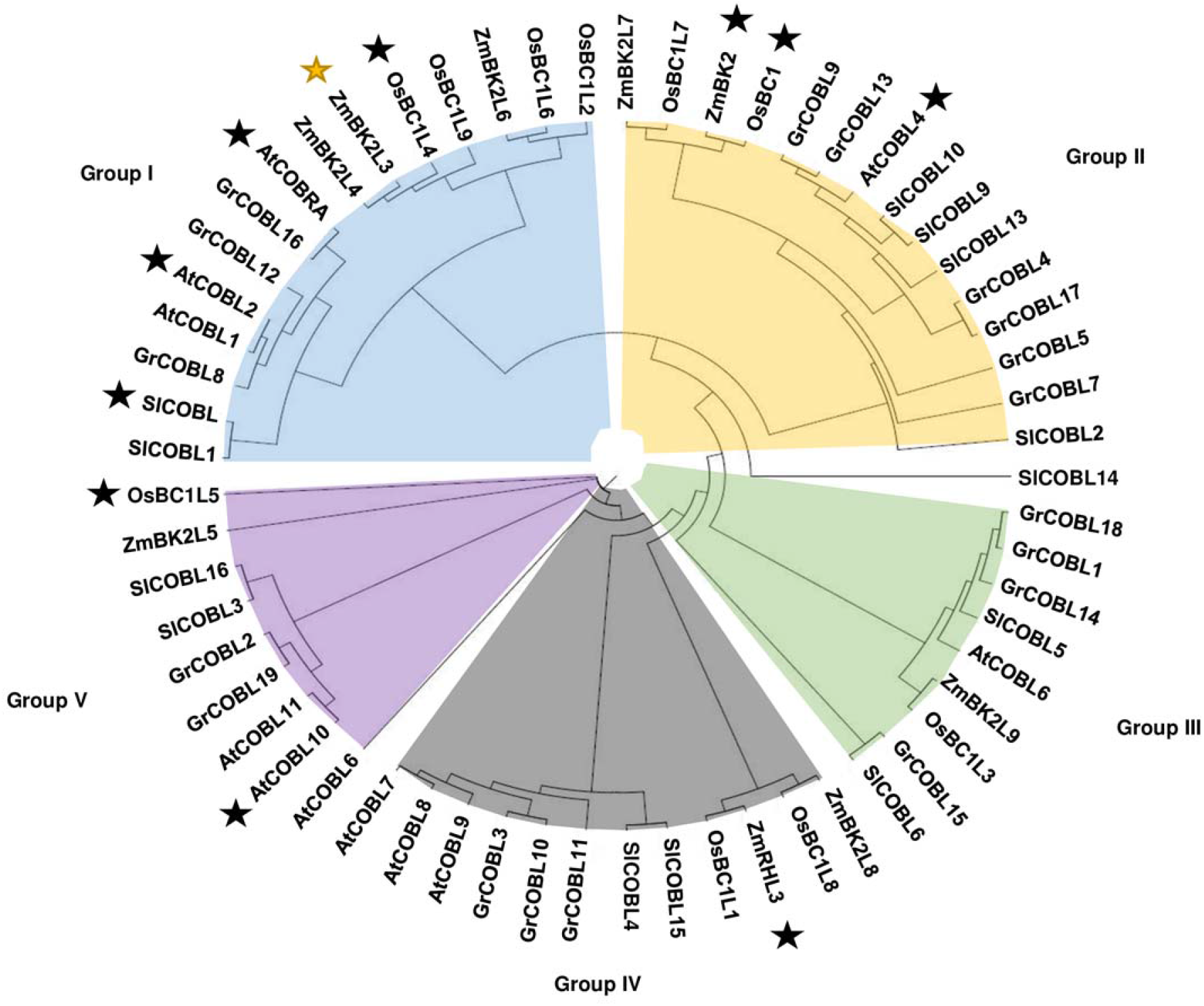
A phylogeny of the COBRA family protein sequences in *Arabidopsis thaliana* (At), *Zea mays* (Zm), *Oryza sativa* (Os), *Solanum lycopersicum* (Sl), and *Gossypium raimondi* (Gr). The COBRA members separate into five distinct clades (groups). ZmBK2L3 is highlighted by a yellow star, whereas black stars represent previously characterized COBRA family members.

### *bk2l3* mutants exhibit cellulose deficiencies and ultrastructural perturbations in mature leaves

To evaluate whether a classical *cobra* mutant cell wall phenotype underlies the observed carbohydrate partitioning-defective phenotype in *cpd28* and *cpd47*, cellulosic and non-cellulosic cell wall sugars in immature sink and mature source leaves of both mutant plants and their respective wild-type siblings were quantified. Cell walls were isolated and digested in 2 M trifluoroacetic acid (TFA) to separate TFA-labile sugar moieties derived from non-cellulosic polysaccharides from acid-resistant cellulose and lignin polymers. Immature sink leaf cell wall composition was broadly similar between mutant and wild-type siblings for both mutants (Figure 7A; Supp. Table S2). However, alterations in cell wall composition were apparent in mature tissues between mutant and wild-type leaves (Figure 7B; Supp. Table S2). A significant reduction in cellulose per unit dry mass was seen for both *cpd28* and *cpd47* mutants relative to wild type (Figure 7B). Interestingly, a stronger reduction was observed for *cpd28* than *cpd47* (17% and 14%, respectively), consistent with the above growth data indicating that *cpd28* is the more severe mutant allele. Minor, but consistent pleiotropic decreases in the amounts of xylose and galactose, and minor increases in arabinose, were found for both mutant tissues, suggesting slight changes to arabinoxylan and pectin composition (Supp. Table S2). These results indicate that whereas developing sink leaves exhibit comparable cellulose levels to wild type, *BK2L3* function is required for normal levels of cellulose deposition in mature source leaves.

**Figure 7.**
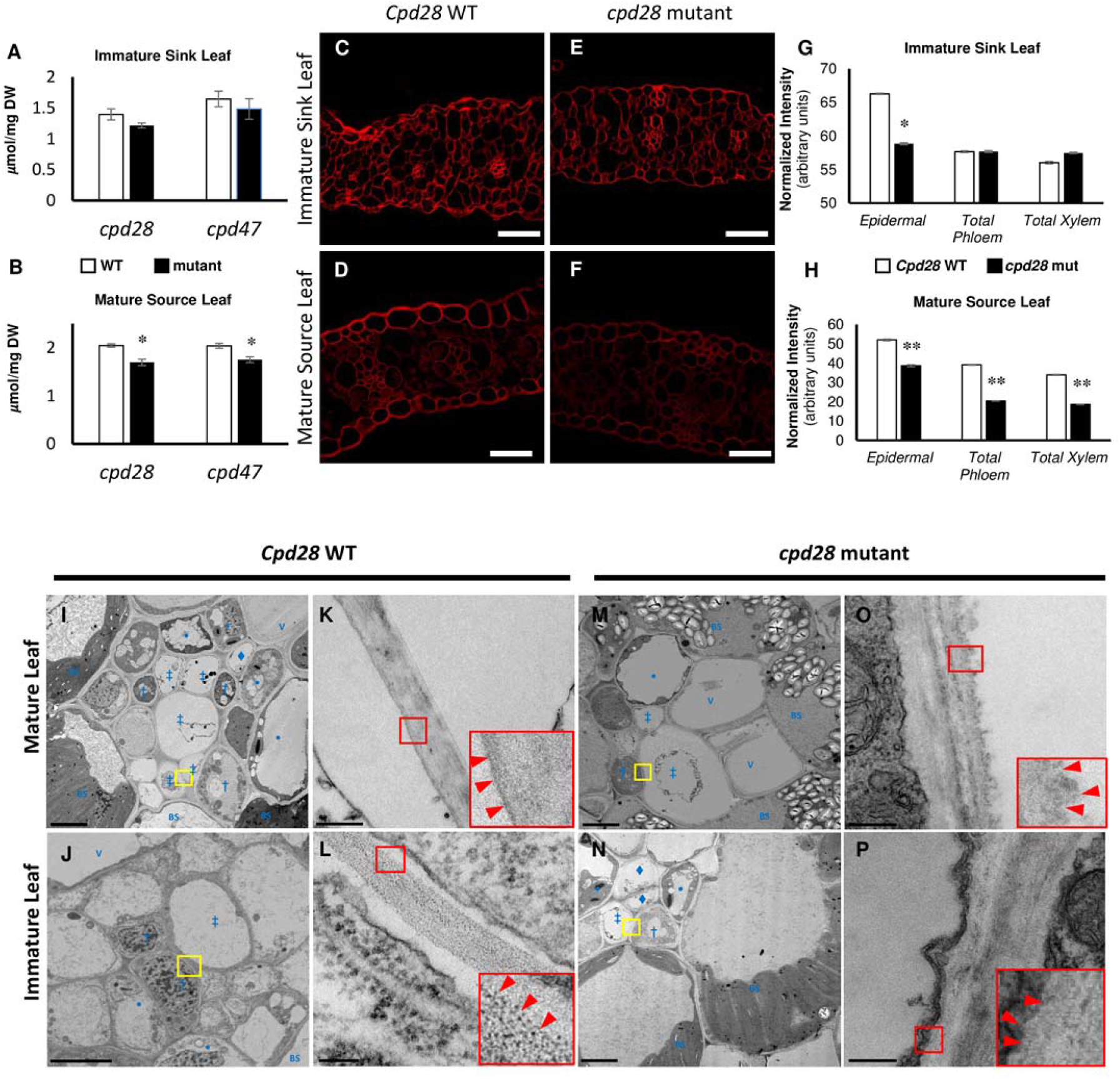
*cpd28* and *cpd47* mutants exhibit a developmentally progressive reduction of cellulose content in comparison to their respective wild-type (WT) siblings. Measurements of cellulose (A, B) in bulk tissue of respective WT siblings, *cpd28*, and *cpd47* mutant immature sink leaves and mature source leaves show reduced cellulose in mature leaf tissue of both mutants. Fluorescence intensity of the cellulose-specific dye Direct Red 23 was used to examine reduced cellulose abundance in mature leaf tissue of the mutant compared to its WT sibling (C-F). Quantification of fluorescence intensity revealed a reduction in cellulose content at the tissue level (G, H). Significant reduction in cellulose abundance was observed in the cell walls of phloem, xylem, and epidermal tissues in mature source leaves (C, E, H) and a small reduction was detected in epidermal tissues of immature sink leaves (D, F, G) of mutants compared to their WT siblings. (I-P) TEM images of veins of mature (I, K, M, O) and immature leaves (J, L, N, P) revealed an abnormal cell wall morphology in thin-walled sieve tubes of mature mutant leaves (M, O), which appeared disorganized and irregular when compared to their WT siblings (I, K). Small red boxes in K, L, O, P show the corresponding magnified inset in the lower right of each panel, with arrowheads highlighting the margin of the sieve tube cell wall. Error bars are mean ± StdEr. *, P ≤ 0.05; **, P ≤ 0.001. Significance calculated using Student’s *t*-test. White scale bar = 50 µm (C-F); black scale bar = 5 µm (I, J, M, N) or 0.2 µm (K, L, O, P). Yellow boxes in lower magnification TEM images (I, J, M, N) highlight the location of the corresponding higher magnification images (K, L, O, P). For I-P cell identities are denoted as BS = bundle sheath, V = xylem vessel, • = vascular parenchyma cell, † = companion cell, ‡ = thin-walled sieve tube, ♦ = thick-walled sieve tube.

To examine cellulose abundance at the cellular level, fixed mature and immature leaves were embedded, sectioned, and stained with Direct Red 23, which binds cellulose and fluoresces red (Thomas et al., 2017). Qualitative inspection of mature tissues showed a marked decrease in red fluorescence in *cpd28* mature leaves compared to wild-type siblings, whereas there were no apparent differences comparing immature leaves from mutant and wild type (Figure 7C-F). To quantify these results, ImageJ analysis of fluorescence intensity of the cell walls from phloem, xylem, and epidermal tissues was performed. In immature *cpd28* mutant leaves, a modest reduction of 11% was observed in cellulose fluorescence of the epidermis, with no significant changes in either phloem or xylem cellulose staining, compared with wild type (Figure 7G). However, in mature leaves, *cpd28* mutants exhibited a ∼26-48% reduction in cellulose staining in all three tissues in comparison with wild type (Figure 7H). These data suggest that BK2L3 is required for normal cellulose deposition in phloem, xylem, and epidermal tissues, and loss of function of the gene significantly reduces cellulose abundance in the vasculature and epidermis of mature leaves.

Sieve element cell walls must resist the high turgor pressure caused by the high osmolarity of phloem sap. Hence, the changes in cell wall composition, in particular the reduction of cellulose in the phloem cell walls, suggest a possible explanation for the carbohydrate export defect in *bk2l3* mutants. To test this hypothesis, we employed transmission electron microscopy (TEM) to examine the ultrastructure of wild-type leaves and *cpd28* mutant tissues exhibiting reduced cellulose. We observed an unusual feature in the cell walls of both the thin-walled sieve tubes (Figure 7M-P) and the thick-walled sieve tubes (Supp. Figure S11A-H) of *cpd28* mutant source leaves, which were not evident in cell walls of corresponding wild-type sieve tubes. While mature wild-type sieve tube cell walls from source leaves appeared smooth at the inner (plasma membrane-facing) side, in *cpd28* mutants, mature sieve tube cells exhibited a ruffled or fuzzy appearance along the plasma membrane interface (Figure 7O, P). Cell walls of sieve tubes of immature wild-type leaves (Figure 7J, L; Supp. Figure S11B, D) appeared smooth and continuous (Evert, 1977), while sieve tubes of *cpd28* mutant immature tissue exhibited a less uniform, grain-like organization, with globular puncta sometimes observable throughout the wall (Figure 7N, P; Supp. Figure S11F, H), which were never observed in corresponding wild-type cell walls. We also observed an irregular cell wall morphology in epidermal cell walls of immature mutant leaf tissue (Supp. Figure S11I, J), including a similar lack of a smooth cell wall surface in the mutant as was observed in the phloem.

### *cpd28* exhibits decreased phloem pressure in mature leaves

The *cpd28* and *cpd47* mutants produce plants with reduced cellulose in mature leaves and altered sieve tube cell wall architecture, which might be responsible for hyperaccumulation non-structural carbohydrates in their leaves because of decreased ability to export photosynthates from source-to-sink. Thus, decreased structural integrity of the sieve element cell wall because of deficiencies in cellulose deposition might compromise the ability of the sieve tube to withstand the high turgor pressure levels needed to drive bulk-transport of photosynthates, as outlined in the pressure-flow hypothesis (Münch, 1930). To test this hypothesis, phloem pressure measurements were performed on *cpd28* mutant and wild-type mature source leaves. The *cpd28* phloem sieve tubes exhibit significantly decreased phloem pressure levels relative to their wild-type siblings (Table 1). Collectively, our data suggest that the impaired carbohydrate partitioning in the *cpd28* and *cpd47* mutants is due, at least partly, to structurally impaired sieve element cell walls with decreased cellulose content being incapable of withstanding the high turgor pressure needed for proper photoassimilate bulk flow from source tissues.

**Table 1.**
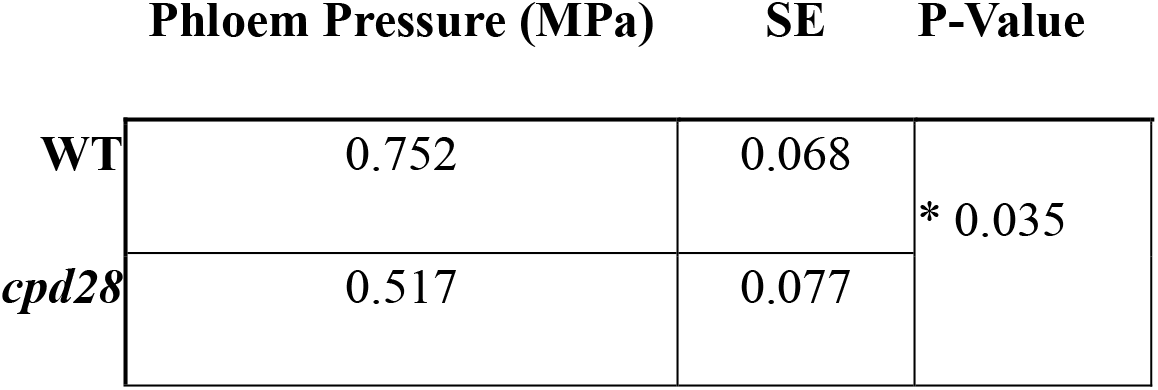
cpd28 mutants exhibit significantly reduced phloem pressure in mature source leaves compared to wild-type (WT) siblings. An approximate 33% reduction was observed. * P ≥ 0.05 by Student’s two-tailed *t-*test. SE, standard error.

## DISCUSSION

The *cpd28* and *cpd47* mutant alleles display a suite of phenotypes similar to other previously characterized mutants inhibited in their carbohydrate transport, such as decreased plant height and vigor, anthocyanin accumulation, and chlorosis of mature source leaves (Lalonde et al., 2003; Baker and Braun, 2008; Braun and Slewinski, 2009; Ayre, 2011). Starch and soluble sugar quantifications and ^14^C-Suc transport studies suggest *cpd28* and *cpd47* mutants hyperaccumulate carbohydrates due to decreased Suc export from the mature source leaves. The observed chlorosis and reduced photosynthetic measurements in the mutant plants likely result from high Glc and/or Suc levels in the leaves, which are known to downregulate photosynthetic gene expression (Rolland et al., 2006; Ruan, 2014). The *cpd28* and *cpd47* mutants also exhibited dwarf phenotypes when grown in dark conditions likely resulting from decreased anisotropic cell elongation, with *cpd28* being more severe than *cpd47*. In addition, *cpd28* mutant mature leaves exhibited a ∼17% reduction in cellulose compared to wild-type siblings, and Direct Red 23 staining also showed that phloem and xylem tissues contained significantly less cellulose in *cpd28* mature leaves compared with wild-type siblings. Consistent with the differences in phenotypic severity, the *cpd28* mutation is a predicted null allele resulting from the loss of the start codon in the *BK2L3* gene. However, the *cpd47* mutation results in a missense mutation leading to an Asp to Asn substitution immediately following the carbohydrate binding motif, resulting in an additional predicted *N*-glycosylation site. COBRA proteins are *N*-glycosylated, and in rice the carbohydrate binding motif is crucial for proper function of the COBRA *BC1* gene (Sato et al., 2010; Liu et al., 2013). Therefore, we suggest that the *cpd47* mutation is a hypomorphic allele.

Previous characterization of other maize mutants which hyperaccumulate non-structural carbohydrates in their leaves identified three different biological processes involved in the export of sucrose from source tissue, namely diminished symplastic transport between companion cells and sieve elements in mature leaves (Braun et al., 2006; Baker and Braun, 2008; Ma et al., 2009; Baker et al., 2013; Tran et al., 2019), reduced apoplastic sucrose phloem loading by SWEET13s and SUT1 (Slewinski et al., 2009; Slewinski et al., 2010; Baker et al., 2016; Bezrutczyk et al., 2018), and ectopic callose formation blocking sieve tubes or specific cell wall interfaces (Russin et al., 1996; Julius et al., 2018). Our characterization of mutants in the *Bk2L3* gene (*cpd28* and *cpd47*) demonstrates that the proper formation of the primary cell wall, in particular cellulose deposition in phloem tissues, is also essential for long-distance carbohydrate partitioning.

Of the currently characterized COBRA genes, none has been reported to have a carbohydrate export defect. However, in addition to the dwarfed phenotype and decreased stem cellulose, rice *osbc1l4* mutants exhibited increased levels of starch in their stems relative to wild-type siblings (Dai et al., 2011). The authors suggested the starch accumulation was due to the inability to incorporate sugars into cell wall components, resulting in the unincorporated Glc being converted to starch. Additionally, in the *Arabidopsis* null mutant *cell elongation defect1* (corresponding to the *Atcobra-5* and *Atcobra-6* alleles) anthocyanin accumulated in aerial tissues, and callose deposition was observed in mutant leaves (Ko et al., 2006). These phenotypes are similar to those observed in *cpd28* and *cpd47* mutants in maize. While there were no reports of increased carbohydrate levels in the *cobra* mutants, defense-related genes were upregulated in *cob*-*5* and *cob-6* compared to wild-type siblings (Ko et al., 2006). In other carbohydrate-accumulating mutants, it was reported that increased carbohydrates might prime defense responses (Gebauer et al., 2017; Julius et al., 2018). Hence, future studies are required to determine if either *Atcobra* or *osbc1l4* mutants accumulate carbohydrates in their leaves when grown under high light conditions.

Members of the COBRA family form five phylogenetic clades, and certain subclades are associated with cellulose biosynthesis during primary or secondary cell wall formation in leaf, stem, root hair, and pollen development (Brady et al., 2007). *Bk2* was hypothesized to function in forming proper cellulose-lignin interactions during secondary cell wall synthesis, and mutations resulted in loss of tissue flexibility, causing brittle plants to snap under slight pressure (Sindhu et al., 2007). The *bk2* plants reach normal height, and no anthocyanin accumulation or chlorosis was observed in the mature leaves (Sindhu et al., 2007). Additionally, *bk2* plants do not hyperaccumulate starch or soluble sugars in their mature source leaves. The *bk2* brittle phenotype is not observed in *cpd28* and *cpd47* mutants. Instead, *cpd28* and *cpd47* plants are dwarfed compared to wild-type siblings. This observation, along with the reduced anisotropic cell expansion in the *cpd28* primary root and reduced expansion of leaf mesophyll cells, suggests that the *BK2L3* gene functions in primary cell wall development in multiple cell and tissue types. In addition, we show here that etiolated *bk2l3* mutants display reduced root and shoot lengths compared to their wild-type counterparts, indicating that like Arabidopsis COBRA, BK2L3 plays a role in rapidly expanding organs. Cellulose orientation is required for anisotropic cell expansion in plants (Daher et al., 2018), therefore, improper formation of primary cell walls would inhibit elongation of growing tissues. This was reported in both primary cell wall-acting *Atcobra* and the *osbc1l4* mutant in rice, which both exhibit dwarf phenotypes and are closely related to *ZmBK2L3* (Li et al., 2003). Additionally, specific *cellulose synthase* genes responsible for primary cell wall growth are co-expressed with *BK2L3* in maize gene expression studies (Dhugga, 2001; Appenzeller et al., 2004; Brady et al., 2007).

The expression profile of *BK2L3* RNA in a developing maize leaf is consistent with expression during both proto- and metaphloem development (Evert et al., 1996; Li et al., 2010; Wang et al., 2014). Thus, the large decrease in cellulose content in mature, fully penetrant source tissue occurs after growth and development have ceased, and is likely a pleiotropic effect similar to that reported in *osbc1l4* (Dai et al., 2011). Therefore, we hypothesize that defects in primary cell wall development in *bk2l3* mutants, particularly in the phloem, result in decreased Suc export, and subsequent build-up of carbohydrates, in the mature leaves.

One hypothesis to link the cell wall defects and growth retardation in the *cpd28* and *cpd47* mutants with the excess carbohydrate accumulation and resulting leaf phenotypes is that it is the decreased sink demand, e.g., from smaller roots, that results in an abundance of carbohydrate in the source tissue. However, if this hypothesis were true, it would be expected that other maize mutants with similarly diminished sink demand, such as *rootless with undetectable meristems1*, *rootless concerning seminal and crown roots1*, *rootless1*, the *defective kernel* and *embryo lethal* classes of mutants, *vanishing tassel2*, and *barren inflorescence2,* to name a few, should present similar carbohydrate accumulation phenotypes in their source tissue, which is not the case (Jenkins, 1930; Neuffer et al., 1997; Woll et al., 2005; McSteen et al., 2007; Taramino et al., 2007; Phillips et al., 2011). Furthermore, the *carbohydrate partitioning defective* phenotypes described in *bk2l3* mutants, as well as others with impaired sucrose export from source tissues (see Introduction), are not observed in mutants with diverse causes for decreased sink demand. Therefore, we suggest that the hypothesis that mutations that result in diminished sink demand result in a carbon hyperaccumulation phenotype in leaves, such as that observed in *cpd28* and *cpd47* mutants, is unlikely.

Alternatively, the source-to-sink transport of carbohydrates, such as Suc, necessitates a particular cellular physiology to accommodate the high pressures accompanying phloem loading. The pressure-flow hypothesis predicts that high concentrations of sugar and other osmolytes increase phloem hydrostatic pressure in source tissues, thus driving the transport of phloem sap to distant tissue sinks (Knoblauch et al., 2016). To compensate for the increased pressure, phloem cell walls must counteract these forces. Interestingly, recent studies using atomic force microscopy (AFM) found that phloem cell walls were significantly more elastic than that of other cell types in *Miscanthus* stems (Torode et al., 2018). Using high resolution AFM to probe sieve element cell walls, the authors observed a spatial distribution of cell wall elasticity, with the highest elasticity observed towards the innermost region of the cell wall. This unique aspect of phloem cell wall physiology may be critical for facilitating high turgor pressures required for long distance phloem transport, and improper formation of the sieve element cell wall is expected to impair long-distance phloem sap transport (Barratt et al., 2011). In the grasses *Triticum aestivum* and *Aegilops comosa*, the innermost, load-bearing layer of both proto- and metaphloem is comprised of dense parallel arrays of radially oriented cellulose microfibrils (Eleftheriou and Tsekos, 1982; Eleftheriou, 1987). If the correct formation of this load-bearing wall were impaired, the cells may not appropriately respond to the high turgor pressure required in the mature source phloem needed to drive bulk flow to sink tissues (Lalonde et al., 2003; Patrick, 2012). Our data show reduced cellulose content in *cpd28* mutant vasculature, and TEM observations of *cpd28* mature and immature mutant leaves indicate that BK2L3 function is required for proper sieve tube cell wall formation, which we interpret as a direct link between primary cell wall formation and Suc export from source leaves via the phloem. Consistent with these observations, phloem pressure measurements of *cpd28* mutant and wild-type mature leaves determined that the mutants exhibited significantly decreased phloem pressure, substantiating the hypothesis that cell wall biophysical properties are critical for maintaining Suc export by bulk-flow. Hence, our data support that BK2L3 is critical for proper cellulose deposition and cell wall formation during phloem development, which is necessary to establish and maintain the high phloem turgor pressure in source leaves.

## MATERIALS and METHODS

### Plant materials and growth conditions

Maize (*Zea mays* L.) plants used for the photosynthetic and gas exchange analyses, microscopy, and soluble sugar and starch quantifications were grown in the field at the University of Missouri South Farm Agricultural Experiment Station in Columbia, MO. The *cpd28* and *cpd47* mutations were generated through pollen EMS mutagenesis of B73 by Patrick Schnable and Nathan Springer, respectively (Liu et al., 2010). Both mutations were backcrossed to the B73 inbred three or more times prior to analyses.

Plants used for the ^14^C-Suc transport assays and microscopy were grown in a greenhouse supplemented with a 1:1 mix of metal halide and high-pressure sodium lighting (1600 μmols m^-2^ s^-1^) under 16-hour light and 8-hour darkness. The plants were kept at a day- and night-time temperature of 30°C and 24°C, respectively. Additionally, etiolated plants used for microscopy and height measurements were grown for 13-15 days under 24-hour dark conditions at 22°C. Genomic DNA was extracted from plants and sequenced to genotype *cpd28* and *cpd47* homozygous mutants, heterozygotes, and homozygous wild-type individuals. A dCAPS assay was developed to allow for easier genotyping of both *cpd28* and *cpd47* mutant alleles. After PCR amplification, DNA from individuals segregating the *cpd28* allele were digested with either *Hae*III or *Alu*I restriction enzymes, where the wild-type allele is cut with *Hae*III and the *cpd28* mutant allele is cut with *Alu*I. Families segregating the *cpd47* allele were PCR amplified and digested with *Hinf*I to cut the wild-type allele. All primer sequences are listed in Supp. Table S1.

*Nicotiana benthamiana* plants used for transient protein expression were grown in a growth chamber under 12 h light and 12 h dark conditions at 18-25**°**C and 55-65% relative humidity for 4-5 weeks (Tran and Braun, 2017).

### Starch staining

Leaves were cleared and stained with IKI as previously described (Baker and Braun, 2007).

### 14C-Suc transport studies

^14^C-Suc was purchased from PerkinElmer (USA) and transport assays were performed on 5 week old plants as described previously (Tran et al., 2019). N = 4 plants were used for *cpd28*, *cpd47*, and wild-type sibling plants, respectively, for each experiment; the experiment was repeated twice.

### Photosynthesis and gas exchange measurements

Gas exchange and photosynthesis measurements were performed on mature source leaves of plants grown in the field between 9:00 AM and 12:00 PM using a portable infrared gas exchange system (LI-6400XT, LI-COR Inc., Lincoln, NE, USA) as described (Huang et al., 2009; Bihmidine et al., 2015). Net photosynthesis (A_net_, μmol CO_2_ m^-2^ s^-1^) and stomatal conductance (g_s_, mol H_2_O m^-2^ s^-1^) rates were measured at a photon flux density of 2,000 μmol m^-2^ s^-1^ and ambient CO_2_ concentration of 400 μmol mol^-1^. N = 5 each for the wild-type, *cpd28*, and *cpd47* plants.

### Soluble sugar and starch quantification

Samples of mature source leaf tissue were harvested from field-grown plants at 5:30 AM and immediately placed in liquid nitrogen before being stored at -80°C until measurement. Soluble sugar and starch samples were extracted according to Leach and Braun (2016). Samples were quantified using high-performance anion exchange chromatography against known standards according to Leach et al. (2017). N = 5 each for *cpd28*, *cpd47*, and their respective wild-type siblings.

### Water relations measurements

Water potentials of leaf tissue were obtained using isopiestic thermocouple psychrometry as described previously (Boyer and Knipling, 1965). Leaf tissue was sampled from mature, pre-anthesis, field grown plants at approximately mid-morning. Osmotic potentials were measured after first measuring water potentials, then freezing tissue overnight at -20C and remeasuring the following day. To obtain turgor pressure values, obtained values were related using the formula Ψ_W_ = Ψ_S_ + Ψ_P_, in effect subtracting the osmotic potential value from the water potential value of a given sample. The experiment was performed twice, and the replicate data sets were determined to be statistically similar by performing a two-way ANOVA, which did not find the individual experiment to be a significant factor using the model Tissue_WP ∼ experiment * sample. Data from the two experiments was then pooled for statistical analysis; with n=7.

### Root growth assay

Seedling root growth was assayed using the growth conditions described (Sharp et al., 1988; Voothuluru et al., 2016). Briefly, 100-150 segregating seeds from a self-fertilized heterozygote were surface-sterilized in 1% NaClO for 15 min, rinsed under running water for 20 min, and treated with 0.1% myclobutanil (Chemsico Inc., St. Louis, MO, USA) for 20 min. Seeds were imbibed for 24 h in 1 mM CaSO_4_ at room temperature with constant aeration, and germinated for 48 h on germination paper moistened with 1 mM CaSO_4_ at 29°C in darkness. Germinated seedlings with primary root lengths between 0.5-2 cm were sown in Plexiglas boxes containing well-watered vermiculite moistened to drip-point with 1 mM CaSO_4_, and grown for 48 h under non-transpiring conditions at 28°C in darkness. Seedlings were removed from the growth medium, primary root axial lengths were measured, and shoot tissue was harvested for genotyping. Seedling roots grew vertically without contacting the bottom of the container for the duration of the growth experiment. The tissue was fixed, embedded, and sectioned as described below for the Direct Red 23 studies. The sections were stained with 0.1% Toluidine Blue O (TBO). N = 19 and 7 for wild-type and *cpd28* roots, respectively. N = 48 and 20 for wild-type and *cpd47* roots, respectively.

The mobilization of carbohydrate from germinating kernels was investigated by assaying root growth on large plates (245 x 245 x 25mm) containing ½ strength MS media and 2% phytagel (Sigma Aldrich #P8169), pH buffered to 5.7 with or without the addition of 2% Suc. Seeds were sterilized and imbibed as with the vermiculite experiments using sterile solutions, and germinated for 48 h on sterile germination paper moistened with 1 mM CaSO_4_ at 28°C in darkness. 18-22 germinated seedlings of roughly uniform root lengths were transferred to each plate; the plates were sealed with parafilm and kept near vertical for 48 h in darkness at 28°C before measuring root lengths and genotyping each seedling.

### Root cell number, length, and width measurements

All cell measurements were made using the ImageJ processing and analysis software (imagej.nih.gov/ij/) from composite TIF images of TBO stained longitudinal sections of wild-type and *cpd28* mutant primary root tips taken on a Zeiss Axiovert 200M motorized widefield microscope. Cell lengths and widths were measured in 1 mm intervals of primary root tip longitudinal sections starting at 1 mm from the root cap junction and ending at 11 mm distal. Between 85-160 cells were measured per region per genotype. Due to their small size, the number of cells in the 1-2 mm apical region were counted from 6-9 separate cortical cells files in similar positions of each region of each root. Because of their larger size, from 2–11 mm from the root cap junction, cell numbers in 3 cortical cell files were counted. Averages are the grand mean from cells within like regions of like genotypes. Statistical significance was determined by a single factor ANOVA. N = 8 roots per genotype.

### Mesophyll cell size measurements

Three individual plants (greenhouse grown, V8 stage) from each genotype were measured, with samples collected from developmentally equivalent leaves cut in the same locations on each leaf. The immature tissue was obtained by dissecting the whorl while the transitioning leaf material was sampled at the boundary where an emerging leaf protrudes above the whorl. Fresh issue was hand-sectioned and stained with aniline blue for visualization of cellular boundaries. Images were collected using the microscope described below, and quantified using ImageJ 1.52. Statistical significance was calculated by one-way ANOVA (P ≤ 0.01) with groupings obtained by Tukey-Kramer post-hoc using the R software package.

### Light Microscopy

All epi-fluorescence images for each experiment were taken using the same microscope and camera settings using a Nikon Eclipse 80i epifluorescent microscope equipped with a 100-W mercury bulb and a DXM1200F camera. Mature and etiolated leaf tissue free-hand cross-sections were generated and imaged as previously described (Baker et al., 2016). Callose was visualized under UV excitation by staining mature leaf sections with 0.1% (w/v) aniline blue in 0.15 M potassium phosphate buffer (pH 8.2) for 5 minutes.

For histological studies of cell walls, field grown mutant and wild-type plants were harvested at the V7 stage and tissue was collected from mature leaves and immature leaves dissected from within the whorl. Samples were prepared as described (Ruzin, 1999); briefly, 2 cm x 2 cm sections of fresh tissue were fixed in a 4% paraformaldehyde solution overnight, followed by dehydration in a graded ethanol series. Tissue was embedded in paraffin blocks and 7 μm sections were cut on a Leica RM2135 microtome. For the cellulose quantification imaging, all slides were processed simultaneously and treated identically. Dewaxed slides were stained in 0.2% (w/v) solution of Direct Red 23 in PBS (equivalent to pontamine fast scarlet 4b, Sigma Aldrich catalog number 212490) (Thomas et al., 2017; Aleamotu’a et al., 2018). Slides were stained for 2 minutes, followed by two 20-minute washes in PBS. Slides were mounted in PBS and sealed before imaging. A Leica TCS SP8 confocal microscope system (Leica Microsystems, Germany) was used to collect confocal images, using 561 nm excitation and detection between 580–615 nm with a 60x water immersion objective. For all collected images, microscope settings used were identical; pinhole was held at 0.75 AU throughout and laser power was kept at 30%. Fluorescence signal was quantified using ImageJ 1.52. Fluorescence intensity values were obtained by selecting individual pixels from the center of the cell wall; at least 100 such individual measurements were obtained for any one tissue measurement (see Supp. Table S3 and Supp. Figure S10). Values were normalized by subtracting the mean value of background fluorescence from that same image, which was always close to zero. Data was analyzed for normality and statistical significance was calculated by one-way ANOVA using the R software package.

### Electron Microscopy

Tissue samples were collected from greenhouse grown wild-type and *cpd28* mutant siblings at approximately the V10 stage. Mature leaf samples were collected at mid-leaf from the youngest fully expanded leaves. The immature leaf samples were collected from the partially emerged leaves 15 and 16, which were dissected from the whorl. The sampled immature leaves had a mean length of 39.4 ± 8.8 cm emerged from the whorl, and 25.3 ± 4 cm not yet emerged from the whorl. Immature leaf samples were collected at the approximate midway portion of the leaf still unemerged from the whorl, 11.5 cm from the base of the leaf. Once cut, tissue samples were immediately processed for TEM. Unless otherwise stated, all reagents were purchase from Electron Microscopy Sciences and all specimen preparation was performed at the Electron Microscopy Core Facility, University of Missouri. Tissues were fixed in 2 % paraformaldehyde, 2 % glutaraldehyde in 100 mM sodium cacodylate buffer pH=7.35. Next, fixed tissues were rinsed with 100 mM sodium cacodylate buffer, pH 7.35 containing 130 mM Suc. Secondary fixation was performed using 1 % osmium tetroxide (Ted Pella, Inc. Redding, California) in cacodylate buffer using a Pelco Biowave (Ted Pella, Inc. Redding, California) operated at 100 Watts for 1 minute. Specimens were next incubated at 4 °C for 1 hour, then rinsed with cacodylate buffer and further with distilled water. En bloc staining was performed using 1% aqueous uranyl acetate and incubated at 4°C overnight, then rinsed with distilled water. A graded dehydration series was performed using ethanol, transitioned into acetone, and dehydrated tissues were then infiltrated with a 1v/1v of Epon and Spurr resin for 24 hours at room temperature and polymerized at 60 °C overnight. Sections were cut to a thickness of 75 nm using an ultramicrotome (Ultracut UCT, Leica Microsystems, Germany) and a diamond knife (Diatome, Hatfield PA). Thin sections were stained using Reynold’s Lead Citrate. Images were acquired with a JEOL JEM 1400 transmission electron microscope (JEOL, Peabody, MA) at 80 kV on a Gatan Ultrascan 1000 CCD (Gatan, Inc, Pleasanton, CA).

### Subcellular localization assays

The coding sequence for the pH insensitive Citrine fluorescent protein was inserted in the coding sequence of the BK2L3 protein (Zm00001d034049) immediately after the N-terminal secretory peptide domain and synthesized in the pDONOR221 vector (Life Technologies, USA) (Tian et al., 2004). The BK2L3-Citrine construct was recombined into the pEarleyGate100 vector generating the 35Spro:BK2L3-Citrine translation fusion protein. *Agrobacterium tumefaciens* transformation and tobacco leaf infiltration was performed as described (Tran et al., 2019). A Leica TCS SP8 confocal microscope system was used for imaging. All microscope and camera settings used were identical across samples within an experiment and were performed as previously described (Baker et al., 2016).

### Mapping and whole genome sequencing

Mapping population generation, bulked segregant analysis (BSA), fine mapping, and whole genome sequencing was performed as described previously (Settles et al., 2014; Tran et al., 2019). To identify the mutations, the sequences were aligned to the B73 reference genome version 4. The fine mapping population consisted of 127 mutant and 298 wild-type plants. Sequences of mapping markers can be found in Supp. Table S1.

### Phylogenetic Analysis

A phylogenetic relationship was constructed of the COBRA protein families from *Arabidopsis thaliana*, *Zea mays*, *Oryza sativa*, *Solanum lycopersicum*, and *Gossypium raimondi*. Accessions were obtained through BLASTP in Phytozome using the Arabidopsis COBRA proteins. The settings for the BLASTP analysis were as follows: an E threshold of 0.001, BLOSUM62 comparison matrix, and gaps allowed. Once the COBRA family members were identified for each taxa, an alignment was generated using default parameters with MUSCLE (Edgar 2004). The output FASTA alignment was curated using Gblocks to exclude non-informative sites using the default parameters for protein alignments (Castresana 2000). The resulting curated FASTA alignment was submitted into the maximum likelihood method PhyML to generate the phylogeny in MEGA6 (Tamura et al., 2013). Statistical analysis was based on the approximate likelihood-ratio test (aLRT) (Guindon et al., 2010). In visualizing the final unrooted phylogenetic tree, the placement of proteins reflects their sequence similarity, with the lengths of the branches arranged for maximal readability.

### Cell wall isolation and compositional analyses

Cell walls were isolated from mutant and wild-type sibling plants as described (Mertz et al., 2012). Briefly, 200-500 mg of frozen leaf tissue from field-grown immature sink and mature source leaves harvested at 5:30 AM were weighed, ground with a mortar and pestle under liquid nitrogen, and stored at -80°C until all samples were processed. Frozen powder was suspended in ∼10 mL of 50 mM Tris-HCl, pH 7.2 containing 1% v/v SDS, and homogenized in a 30 mL glass-glass Duall tissue homogenizer (DWK Life Sciences, Rockwood, TN). The homogenate was pelleted at 2000 g for 5 minutes and the supernatant was replaced with fresh buffer and heated to 65°C for 20 minutes to extract soluble proteins. Walls were pelleted and washed 3x each with warm (50°C) dH_2_O, warm 50% ethanol, and room temperature dH_2_O. Isolated walls were resuspended in dH_2_O with 1-2 drops of toluene to prevent microbial growth and stored at 4°C. N = 6 each for *cpd28*, *cpd47*, and their respective wild-type siblings.

For quantification of TFA-labile cell wall sugars, 1 mL aliquots of wall suspension were treated with 2 U porcine α-amylase/mg carbohydrate in 10 mM Tris-Malate, pH 6.9 as described in Pettolino et al., (2012) to remove starch that would confound the analysis of cell wall Glc moieties. 1-2 mg of destarched, lyophilized wall material were hydrolyzed in 2 M TFA containing 500 nmol/μL *myo*-inositol for 90 minutes at 120°C, reduced with NaBH_4_, and converted to alditol acetates as described (Gibeaut and Carpita, 1991). Derivatized sugars were separated and quantified by gas chromatography-mass spectrometry using the parameters described (Mertz et al., 2012).

For quantification of cellulose by the phenol-sulfuric assay, TFA-insoluble pellets of 4-6 mg lyophilized wall material were recovered following hydrolysis and washed 3x with dH_2_O in a 5 mL conical Reacti-Vial (Pierce). Triplicate 200 μL aliquots of suspended pellets from each sample were combined with 5% (v/v) phenol and concentrated sulfuric acid, developed for 2 h, and analyzed spectrophotometrically for Glc equivalents at 500, 510, and 520 nm according to Dubois et al., (1956).

### Phloem pressure measurements

Pico gauge pressure measurements (Knoblauch et al., 2014) were performed on a Leica DM-LFSA equipped with a Leica DC450 digital camera. For in situ phloem pressure measurements, a longitudinal section was removed from the bottom of the main vein (midrib) of a leaf still attached to an intact 8-week-old plant to expose the phloem without damaging it (Supp. Figure S12). This leaf was mounted on the stage of the microscope on a tilted stage (∼20% tilt). Images were taken with a Leica 63x water immersion lens. Image analysis and pressure calculations were performed as described in detail in Knoblauch et al. (2014). N = 8 for *cpd28* mutants and 9 for wild-type siblings.

## Supplemental Data Files

**Supplemental Figure S1.** *cpd47* mutant plants are dwarfed relative to wild-type siblings and exhibit basipetal accumulation of anthocyanin in their leaves.

**Supplemental Figure S2.** *cpd28* and *cpd47* mature leaves exhibit decreased photosynthesis and stomatal conductance.

**Supplemental Figure S3.** *cpd47* mutants exhibit reduced Suc export from source tissues leading to hyperaccumulation of starch and soluble sugars in mature leaves.

**Supplemental Figure S4.** Aniline blue staining of wild-type, *cpd28* and *cpd47* mutants leaf cross sections.

**Supplemental Figure S5.** Supplementing growth media with Suc did not rescue the reduced root elongation phenotype of *cpd28* mutants.

**Supplemental Figure S6.** Mesophyll cells exhibit reduced expansion in *cpd28* mutant leaves.

**Supplemental Figure S7.** The *cpd47* mutation occurs at the end of the cellulose binding motif.

**Supplemental Figure S8.** Soluble sugar and starch quantification in wild-type and *bk2* mature leaves.

**Supplemental Figure S9.** *BK2L3* is most highly expressed in expanding tissues.

**Supplemental Figure S10.** Workflow for quantification of Direct Red 23 fluorescence.

**Supplemental Figure S11.** TEM images of thick-walled sieve tube and epidermal cell walls.

**Supplemental Figure S12.** Experimental set up for phloem pressure measurements.

## ACKNOWLEDGMENTS

We appreciate the insightful comments of three anonymous reviewers and the editor for suggestions that greatly improved the manuscript. The authors would like to thank Trupti Joshi for help with whole genome sequencing data analysis. We would also like to thank the MU Molecular Cytology Core for their advice and expertise in confocal microscopy. We would also like to thank George Chuck and David Collings for sharing their Direct Red 23 staining protocols and Anna Olek for laboratory assistance with the cell wall analyses. No conflicts of interest declared.

## Author Contributions

B.T.J. conducted the light and fluorescence microscopy, starch staining, shoot and root length measurement, isolated the gene, drafted the manuscript, and helped critically revise it; T.J.M. conducted the photosynthesis and gas exchange measurements, root length measurements, sucrose-feeding root growth experiments, ^14^C-sucrose transport studies, Direct Red 23 cellulose imaging and quantification experiments, light microscopy, subcellular localization assays, electron microscopy experiments, drafted the manuscript, and helped critically revise it; R.A.M. conducted the cell wall analysis, root length measurements, electron microscopy experiments, drafted the manuscript, and helped critically revise it; N.B. performed the root cell number, length, and width quantitative measurements; K.C. performed the starch and soluble sugar quantification, and fine-mapping; D.A.G conducted electron microscopy sample preparation and imaging; J.K and M.K. performed the phloem pressure measurements; S.B., P.C., J.W., R.W., J.P., K.G. and T.L.S. conducted the fine-mapping; M.M. and N.C. participated in the design of the cell wall analyses, and helped critically revise the manuscript; D.M.B. participated in the design of the study, helped draft, critically revise the manuscript, and agrees to serve as the author responsible for contact and ensures communication.

**Supplemental Figure S1.** *cpd47* mutant plants are dwarfed relative to wild-type (WT) siblings, and exhibit basipetal accumulation of anthocyanin in their leaves. (A) Field grown WT (left) and *cpd47* mutant (right) plants. (B-C) A gradient of *cpd47* mutant (B) and WT (C) leaves from oldest (left) to youngest (right). The mutant leaves accumulate anthocyanins in a basipetal fashion when exposed to sunlight. Supports Figure 1.

**Supplemental Figure S2.** *cpd28* and *cpd47* mutants mature leaves exhibit decreased photosynthesis and stomatal conductance compared to their respective wild-type (WT) siblings. (A) Net photosynthetic rate measured in *cpd28* and *cpd47* mutants (black bars) and their WT siblings (white bars). (B) Stomatal conductance measured in *cpd28* and *cpd47* mutants (black bars) and WT siblings (white bars). Error bars are mean ± StdEr. Asterisks represent a significant difference at the P ≤ 0.05 using Student’s *t*-test. Supports Figure 1.

**Supplemental Figure S3.** *cpd47* mutants exhibit reduced Suc export from source tissues leading to hyperaccumulation of starch and soluble sugars in mature leaves. (A-B) Wild-type (WT) sibling (top) and *cpd47* mutant (bottom) mature leaves before (A) and after (B) IKI stain. (C) Soluble sugar and starch quantification in WT and *cpd47* mutant mature leaves. (D-F) Export of ^14^C-Suc is reduced in *cpd47* mutant source leaves. (D) Dried WT (top) and *cpd47* mutant (bottom) leaves. (E) Autoradiograph showing ^14^C-Suc distribution in WT (top) and *cpd47* mutant (bottom) leaves. (F) Quantification of ^14^C-Suc distribution in WT (white bars) and *cpd47* mutant (black bars) leaves. Translocation was measured according to signal intensity at each distance (in cm) from the application site. Error bars are mean ± StdEr, red points indicate individual measurements comprising the mean (C). Asterisks represent a significant difference at the P ≤ 0.05 using Student’s *t*-test. Supports Figure 2.

**Supplemental Figure S4.** Aniline blue staining of wild type (WT) (A,D,G,J), *cpd28* (B,E,H,K) and *cpd47* (C,F,I,L) leaf cross sections. Strongly phenotypic mutant *cpd28* and *cpd47* leaf tips exhibit callose deposits in phloem of mature leaves (B-C), but not in etiolated 2-week old leaves (H,I) or immature light-grown leaves (K,L). Fresh cut tissue sections were stained with aniline blue and observed under UV light with either an epifluoresence microscope with 10X objective (A-C,G-L) or laser-scanning confocal microscope with 60X objective (D-F). White arrowheads indicate position of callose occlusions in phloem. Scale bars = 50 µm (white), 25 µm (gold). Supports Figure 2.

**Supplemental Figure S5.** Supplementing growth media with Suc did not rescue the reduced root elongation phenotype of *cpd28* mutants compared to wild-type (WT) siblings. Error bars are mean ± StdEr. *, P ≤ 0.05; **, P ≤ 0.001. Significance calculated using Student’s *t*-test. Data represents the sum of two experiments, n ≥ 8 replicates per treatment. Supports Figure 3.

**Supplemental Figure S6.** Mesophyll cells exhibit reduced expansion in *cpd28* mutant leaves compared to wild-type (WT) siblings. Significance calculated by one-way ANOVA (P ≤ 0.01) with groupings (letters) obtained by Tukey-Kramer post-hoc. Red numbers indicate the number of cells measured for that value. Supports Figure 3.

**Supplemental Figure S7.** The *cpd47* mutation (yellow) occurs at the end of the cellulose binding domain (green). A multiple sequence alignment was performed on the protein sequences of the COBRA family members in *Zea mays*. Supports Figure 4.

**Supplemental Figure S8.** Soluble sugar and starch quantification in wild-type (WT) siblings and *brittle stalk2* (*bk2)* mature leaves. Error bars are mean ± StdEr, red points indicate individual measurements comprising the mean. Asterisks represent a significant difference at P ≤ 0.05 using Student’s *t*-test. Supports Figure 6.

**Supplemental Figure S9.** *BK2L3* is most highly expressed in expanding tissues. A. RNA expression along a gradient in a transitioning leaf (immature base to left, mature tip to right). Expression of *BK2L3* shown in blue, *BK2L4* in green, and *BK2* in yellow. Note peaks for expression of *BK2L3* and *BK2L4* occur in younger, expanding tissue, corresponding to primary cell wall formation, whereas *BK2* expression peaks later corresponding to secondary cell wall formation. B. *BK2L3* is expressed at higher levels in the root elongation zone compared to the meristematic zone. Supports Figure 6.

**Supplemental Figure S10.** Workflow for quantification of Direct Red 23 fluorescence. (A) Stained tissue sections were imaged on a confocal microscope using identical parameters. (B) Images were converted to 16-bit grayscale using ImageJ. (C) Phloem pole, xylem pole, and epidermal tissues (not shown) were measured separately (white box in B denotes vein shown in C and D). (D) Fluorescence signal intensity was measured using the ImageJ multi-point tool around the cell walls for tissue of interest (xylem in C shown as an example). (E) The obtained values were observed to be normally distributed before conducting the statistical analysis described in Figure 7 and Supplemental Table S3. Supports Figure 7.

**Supplemental Figure S11.** TEM images of wild-type (WT) sibling and *cpd28* mutants thick-walled sieve tube and epidermal cell walls. Black scale bar = 5 µm (A, B, E, F) or 0.2 µm (C, D, G-J). Yellow box in lower magnification TEM images (A, B, E, F) shows the location of the corresponding higher magnification image (C, D, G, H). Smaller red box in higher magnification images (C, D, G-J) shows the corresponding magnified inset in the lower right of each panel, with arrowheads highlighting the margin of the sieve tube wall. Cell identities are denoted as BS = bundle sheath, V = xylem vessel, • = vascular parenchyma cell, † = companion cell, ‡ = thin walled sieve tube, ♦ = thick walled sieve tube. Supports Figure 7.

**Supplemental Figure S12.** Experimental set up for measuring phloem pressure in a maize leaf. (A) A *cpd28* mutant plant oriented to position a mature source leaf on the microscope stage. (B) A small section of overlying abaxial epidermal and subdermal tissue is removed to expose the phloem. (C) A picogauge is gently inserted into a phloem sieve element to measure the pressure. Supports Table 1.

---

**Supplemental Table S1.**
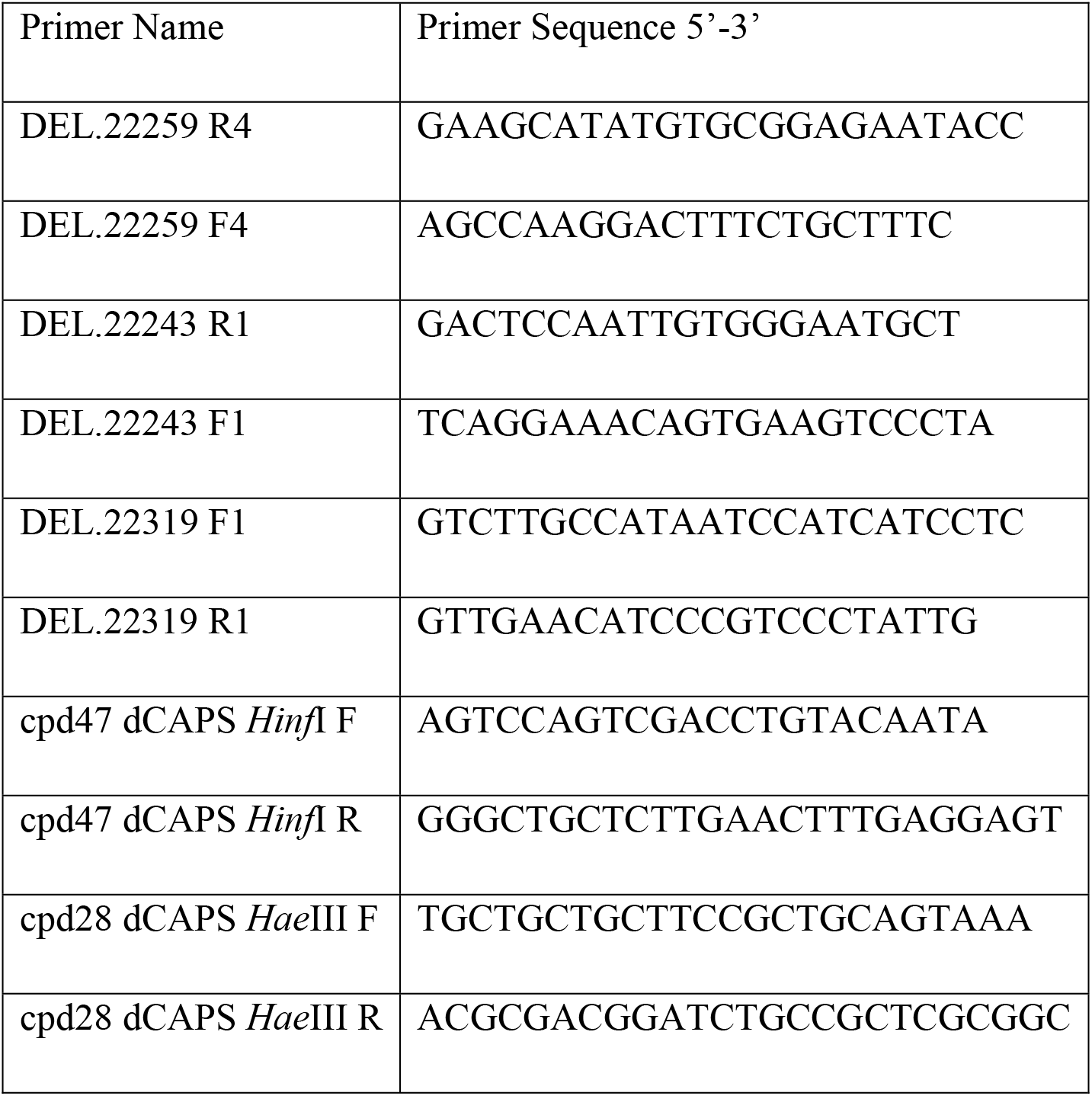
Mapping and genotyping primer sequences

**Supplemental Table S2.** TFA-labile monosaccharides of mutant (mut) and wild-type (WT) cell walls

| **Sugar**<br>**[nmol/mg**<br>**DW]** | ***Cpd28* WT Imm** | ***cpd28* mut Imm** | ***Cpd47* WT Imm** | ***cpd47* mut Imm** | ***Cpd28* WT Mat** | ***cpd28* mut Mat** | ***Cpd47* WT Mat** | ***cpd47* mut Mat** |
| --- | --- | --- | --- | --- | --- | --- | --- | --- |
| **Rha** | 29.3±3.4 | 26.9±2.5 | 21.3±2.6 | 21.5±1.8 | 27.8±1.0 | 26.1±1.5 | 23.6±1.3 | 25.2±0.9 |
| **Fuc** | 6.4±0.6 | 6.8±0.7 | 6.2±0.4 | 7.1±0.6 | 7.9±0.4 | 7.0±0.7 | 6.9±0.4 | 7.5±0.3 |
| **Ara** | 541.7±20.9 | 477.8±0.5 | 497.5±15.9 | 502.5±11.2 | 456.0±14.6 | 498.6±19.3 | 439.1±14.1 | 540.4±14.9 |
| **Xyl** | 1357.6±106.9 | 1233.2±51.4 | 1677.9±54.6 | 1674.5±87.9 | 1430.5±44.8 | 1348.6±23.5 | 1487.2±57.8 | 1476.9±46.6 |
| **Man** | 13.0±2.2 | 13.4±1.4 | 8.1±0.8 | 10.5±0.8 | 7.8±0.3 | 10.3±0.5 | 6.7±0.7 | 7.2±0.7 |
| **Gal** | 118.7±22.6 | 93.2±6.9 | 75.4±7.4 | 70.3±6.8 | 59.1±2.0 | 110.6±11.6 | 52.9±2.0 | 107.7±5.6 |
| **Glc** | 807.4±106.8 | 571.1±36.2 | 601.9±33.6 | 534.8±13.6 | 192.7±5.7 | 215.0±15.3 | 206.4±5.9 | 249.0±10.1 |

Values are means [nmol sugar/mg dry weight] ± StdErr. Imm, Immature; Mat, Mature; Rha, Rhamnose; Fuc, Fucose; Ara, Arabinose; Xyl,
Xylose; Man, Mannose; Gal, Galactose; Glc, Glucose

**Supplemental Table S3.**
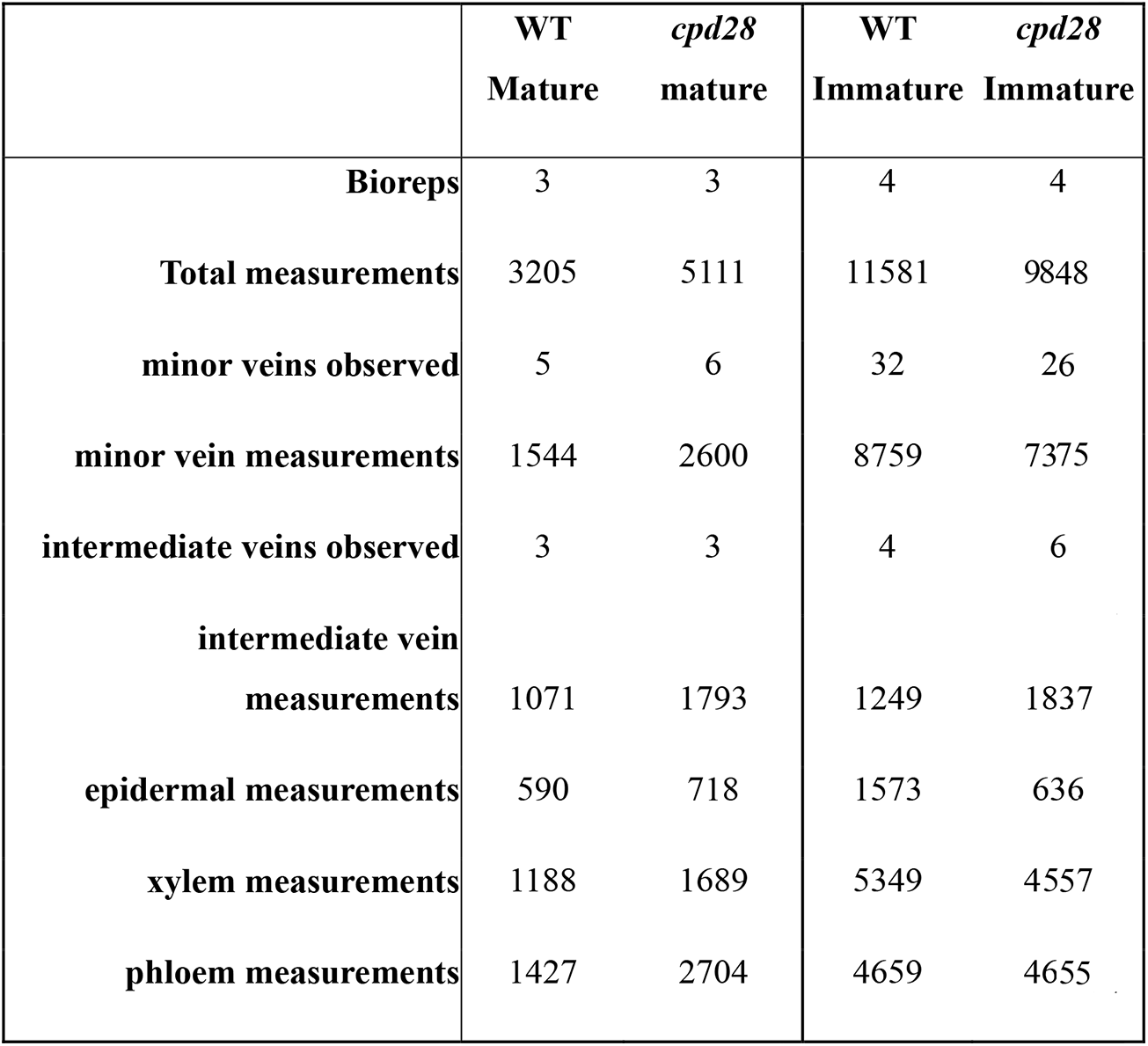
Meta data reporting the observations used for quantification of fluorescence signal as seen in Fig. 7.

|  | WT<br>Mature | *cpd28*<br>mature | WT<br>Immature | *cpd28*<br>Immature |
| --- | --- | --- | --- | --- |
| **Bioreps** | 3 | 3 | 4 | 4 |
| **Total measurements** | 3205 | 5111 | 11581 | 9848 |
| **minor veins observed** | 5 | 6 | 32 | 26 |
| **minor vein measurements** | 1544 | 2600 | 8759 | 7375 |
| **intermediate veins observed** | 3 | 3 | 4 | 6 |
| **intermediate vein**<br>**measurements** | 1071 | 1793 | 1249 | 1837 |
| **epidermal measurements** | 590 | 718 | 1573 | 636 |
| **xylem measurements** | 1188 | 1689 | 5349 | 4557 |
| **phloem measurements** | 1427 | 2704 | 4659 | 4655 |

